# Investigating multiple types of resistance against a homing gene drive in European populations of *Drosophila melanogaster*

**DOI:** 10.1101/2025.06.06.658298

**Authors:** Nicky R. Faber, Jackson Champer, Bart A. Pannebakker, Bas J. Zwaan, Joost van den Heuvel

## Abstract

Gene drive technology may be a valuable tool for addressing several contemporary challenges, including combating disease vectors, conserving biodiversity, and controlling agricultural pests. Homing gene drives spread through a population by copying themselves onto the homologous chromosome in the germline of heterozygous individuals. However, it is possible that resistance will evolve against homing gene drives, especially if the goal is to suppress or eliminate a pest population so that resistance alleles have a large selective advantage over the gene drive. Resistance can result from a simple mutation at the drive’s target site, which is found in many studies but can potentially be avoided by improving drive design. However, a more complex polygenic type of resistance could also evolve through selection on standing genetic variation that affects the efficiency of the spread of the gene drive. In this study, we test an efficient homing gene drive in genetically diverse lines of *Drosophila melanogaster*, collected from across Europe by the DrosEU Consortium. We find that the gene drive shows considerable variability in homing efficiency, but that none of this variability can be ascribed to heritable genetic effects. Selection for complex resistance is thus unlikely and will be inefficient, probably still giving a gene drive enough time to fixate in the population. However, although our tested gene drive targets a highly conserved haploinsufficient gene with two gRNAs, we find simple resistance alleles in viable offspring. Half of these are the product of end-joining repair instead of homing and may still carry heavy fitness costs. However, the other half are the result of partial homing events. These alleles indicate that resistance could likely evolve against this gene drive in a simple, non-polygenic way. Therefore, more effective strategies may be required to address simple resistance mutations, whereas complex resistance may be unlikely to pose a substantial barrier to the employment of at least certain types of gene drive.

## Introduction

Gene drive technology could be a valuable tool for addressing several contemporary challenges, including combating disease vectors such as malaria mosquitoes, as well as conserving biodiversity and managing agricultural pests by controlling invasive species (1). A gene drive is a genetic element that, once released into a population, is inherited at super-Mendelian rates, thereby increasing its frequency over successive generations (2, 3). Synthetic gene drive systems can be designed with two purposes in mind: modification drives, which aim to spread a neutral or beneficial trait, and suppression drives, which purposefully cause a deleterious effect to shrink or eradicate populations of harmful species (4). Although the technology holds significant potential, much work remains to be done before it can feasibly be used for practical applications (5–8). This work includes achieving the necessary high inheritance rates and high fitness of drive carriers for the gene drive to establish and spread in a population (6, 9–11), avoiding the formation of resistance alleles that block the drive and can negate its effects (12–15), achieving localization for global biosecurity (16–19), as well as establishing regulatory frameworks for the governance of this technology, alongside engaging the general public (20, 21).

The evolution of resistance presents a significant obstacle, because it undermines the gene drive’s intended purpose either partially or entirely (22). Especially in the case of suppression drives, resistance is likely to pose a problem due to the large selective advantage that resistant individuals will have once the population is reduced in size. Resistance can take three general forms (22), the first of which is through mutations in the gene drive’s target site, which have been found in many experimental demonstrations of gene drive constructs (12–15). We call this type of resistance “simple resistance” (Figure 1). The mutations at the target site usually arise through the drive’s activity itself (13). However, they can also already be present in a population, and requires screening of the target population beforehand (23–25). This type of simple resistance seems largely avoidable by good gene drive design, such as targeting conserved sequences and gRNA multiplexing (25–32). The other two ways in which resistance could evolve as described by Price et al. (22) are through molecular interference from non-target genetic elements, and through behavioural or life history changes within the population. As these types of resistance are likely to be complex, quantitative traits, we call them “complex resistance” hereafter (Figure 1).

**Figure 1.**
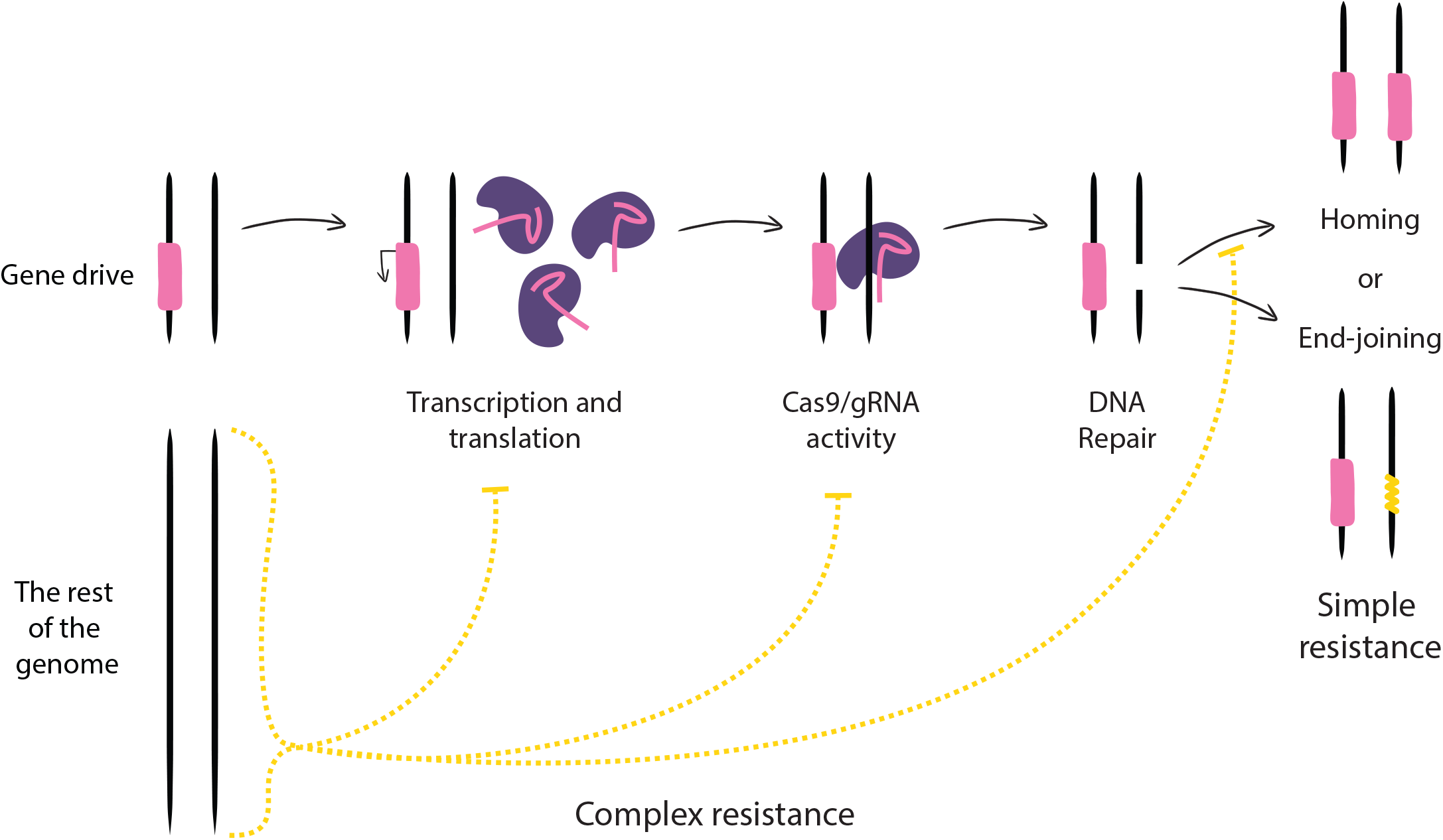
Schematic of simple and complex resistance against a CRISPR-based homing gene drive. Pink and purple show the gene drive components going through the various steps of activity, with the outcome being either homing or end-joining. Yellow lines represent resistance against the gene drive, with the squiggly yellow line representing simple resistance after end-joining repair (instead of homology-directed repair for homing), and the arrows inhibiting different processes indicating complex resistance. We show three processes essential for homing in which complex genomic resistance could theoretically interfere, but there could be many others.

The forms of complex resistance have remained mostly the-oretical (33) and most gene drives are only tested in single lab-oratory strains, such as the white-eyed *Drosophila melanogaster* strain *w*^*1118*^ (34–37). Although there have been a few experimental studies testing a gene drive in different genetic backgrounds, none of them have formally calculated the heritability of their efficiency, nor accounted for confounding factors such as maternal, environmental, or random effects (38–43). Broadly speaking, evolutionary history has demonstrated that resistance can arise against many ecological interventions when there is a significant enough benefit to it (44–50). Also, in theory, there are many molecular processes essential for gene drive functioning for which standing genetic variation may be present in the targeted population and that could convey (partial) complex resistance. In the case of CRISPR-based homing gene drives, which we focus on in this study, Cas9 cutting efficiency could be reduced, DNA repair could switch from homology-directed repair to endjoining, the timing of the expression of the drive may be shifted, the deleterious effect that the gene drive causes could be mitigated, or the recessive genetic load that the drive imposes may not stay fully recessive, among many other potential mechanisms. Thus, anything the gene drive field is currently trying to optimize in their constructs could be counteracted by the evolution of complex resistance (6, 7). Exploring the likelihood that complex forms of resistance arise is important for estimating the evolutionary hurdles that gene drive technology may yet face, especially as some drive constructs may be ready for field trials in the near future (51–53).

Besides simple and complex resistance, there is an additional type of resistance to consider for a specific type of homing gene drive: the homing rescue drive. Some modification gene drives (which aim to drive a trait into a population, instead of to suppress it) are located within a gene while also containing a recoded “rescue” element for this gene. The rescue element needs to be recoded with synonymous substitutions to avoid cleavage by the gene drive’s gRNAs. This construction allows the gene drive to avoid simple resistance by targeting highly conserved genes, because most simple resistance alleles will be deleterious, whereas the gene drive with the rescue element will be fully functional. However, it also puts the gene drive at risk of partial homing, where homology-directed repair could, after copying the rescue of the gene drive, revert back to the wild-type strand without copying the other components of the gene drive. The recoded rescue element still has some homology with the wild-type sequence (usually including part of the 3’ untranslated region of the gene as well), which could facilitate this partial homing. The result will essentially be a simple resistance allele, only it is more likely to be a functional resistance allele than end-joining mutations (which are often out-of-frame). However, this mechanism has not yet been demonstrated in practice for any homing rescue gene drive (54–58).

In this study, we test all these different types of resistance that could evolve against a homing rescue gene drive (54). We investigate the impact of standing genetic variation on gene drive efficiency to estimate the likelihood that complex resistance could pose a problem for gene drives in genetically diverse populations in the field. To this end, we test the gene drive in a broad range of genetically diverse *Drosophilamelanogaster* populations. These populations were sampled by the DrosEU consortium from across Europe (59), and isofemale lines were created as a snapshot that captures the genetic diversity in these populations at the time (60). Specifically, we measure the two main phenotypes indicative of gene drive success to cast as wide a net as possible: gene drive inheritance rate and number of offspring. We find that the gene drive shows considerable variability in inheritance rates, but this variability cannot be ascribed to heritable genetic effects. Also, there is a notable shift in the number of offspring produced between gene drive and control crosses, but again, this shift cannot be linked to the genetic background of the cross. Finally, from crosses that exhibited especially low efficiency, we screened viable offspring for resistance alleles at the gRNA target sites, and found both deletions and partial homing events. We conclude that simple resistance is more likely to evolve than complex resistance, and we suggest that the modularity of gene drive technology can be harnessed to manage this.

## Methods

### A. Nested full-sibling crosses to estimate heritability

To calculate the heritability of gene drive efficiency, we use a nested full-sibling experimental design. We have three levels of nesting, namely population, isofemale line, and sibling batch (Figure 2A). We tested six populations, five isofemale lines from each population, and five sibling batches per isofemale line, with five gene drive males from each batch as biological replicates (full siblings). We only test the heritability of homing efficiency in male drive individuals, because our chosen Cas9 is predominantly active in the male germline. We use a Generalized Linear Mixed Effect model to partition the observed variance to each level in our experiment (61). We consider population-level and isofemale-level effects as genetic effects, whereas the sibling batch is used to correct for potential maternal or environmental effects, and between-sibling variance indicates stochastic effects. We note that there may still be genetic variation within each isofemale line, as they were not inbred upon collection. Therefore, our heritability estimate will be conservative. We calculate the heritability as follows:

**Figure 2.**
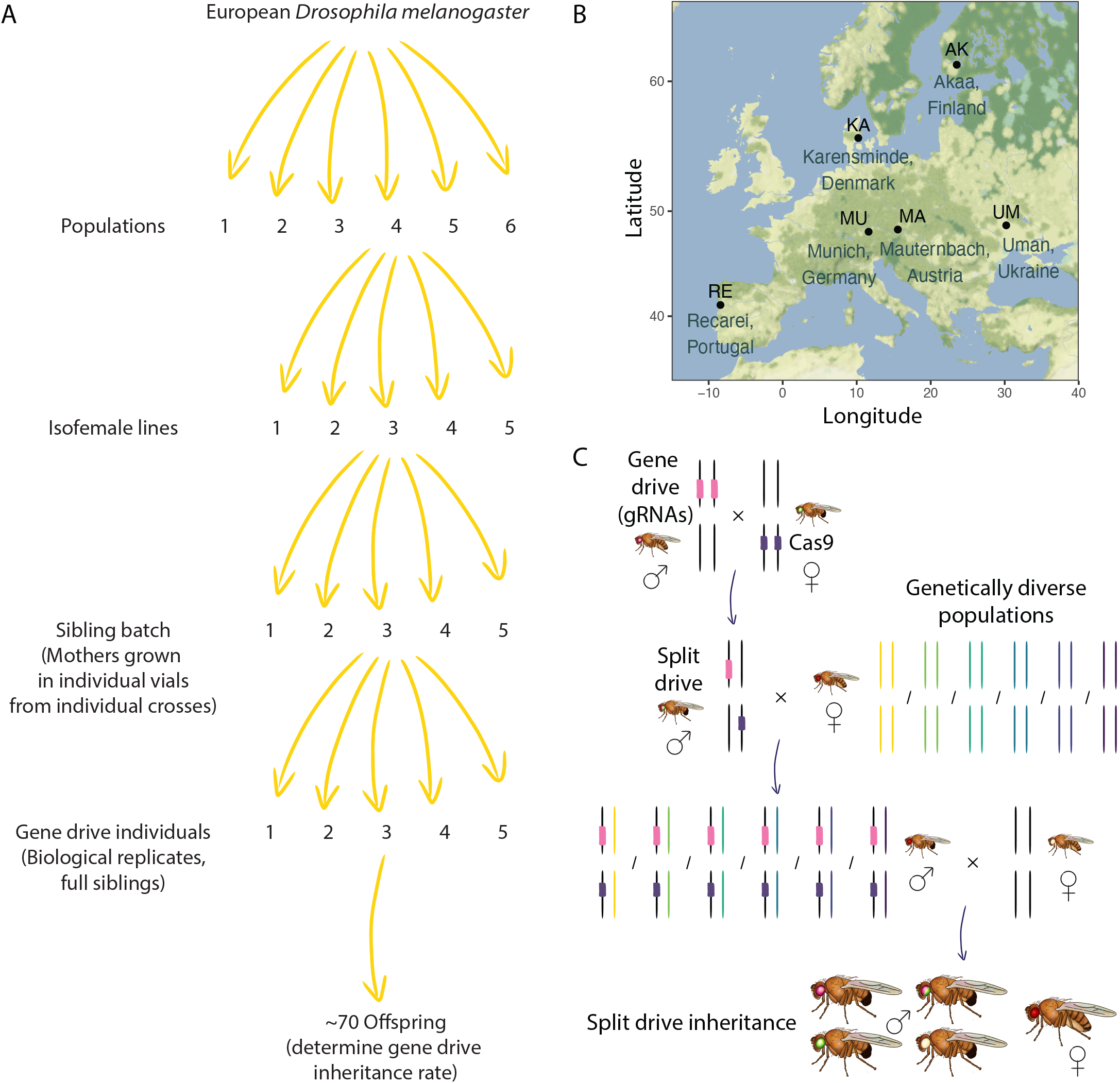
Experimental design to determine the heritability of gene drive efficiency. **A)** We use a nested hierarchical experimental design with four levels: population, isofemale line, sibling batch, and finally, gene drive individual. For illustrative purposes, arrows for the next level down are only drawn for one replicate. **B)** Geographic locations of the six (out of nine in total) *D. melanogaster* populations that were sampled by the DrosEU Consortium and used in this study. **C)** A simplified version of our split homing rescue gene drive crossing scheme.

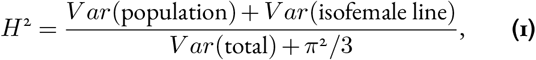

where *π*^2^*/*3 is the variance of binomially distributed data (62).

### B. Choice of genetically diverse *Drosophila melanogaster* lines

The genetically diverse populations we tested were sampled from across Europe by the DrosEU Consortium (59). This consortium sampled from nine populations up to 20 isofemale lines each. We chose six populations from across the East and West populations in which European *D. melanogaster* populations are genetically structured to maximize the amount of genetic variation covered by our experiment (Figure 2B) (59). Among all DrosEU populations, the pairwise *F*_*ST*_ ranges between 0.013 and 0.059, so our chosen populations cover the higher end of that range (59). These lines were kept in laboratory conditions for several years, and some adaptation to the lab conditions is expected. While this generally deemed undesirable to get a realistic measurement of a trait relevant in the field (60), here we are interested in estimating the heritability of gene drive efficiency in general and not so much in whether the underlying genetic variation closely resembles that of the natural source populations.

### C. Choice of gene drive and Cas9

For this study, we used a homing rescue modification gene drive previously built by Champer et al. (54). This gene drive is highly efficient and able to quickly spread through a cage population of *D. melanogaster*. The gene drive is located in the highly conserved haplolethal essential gene *RpL35A*, which encodes for a ribosomal subunit that is expressed at every life stage (63) and in every tissue (64) of *D. melanogaster*. The gene drive targets exon 4 of *RpL35A* gene with two gRNAs, while also, after inserting itself at this location, providing a rescue for the rest of the gene, thus making it a modification gene drive. It should be noted that *RpL35A* is a “*Minute*” locus, meaning that it is haploinsufficient (65, 66), and is presumed to be a haplolethal gene (54). Therefore, most or all simple resistance alleles, either from the germline itself or in the embryo from maternal deposition of Cas9 and gRNAs, result in non-viable offspring as there will be still one malfunctioning copy of *RpL35A* present after the mutational event.

This drive is also a split drive for the molecular safeguarding of experiments (67, 68), which gave us the opportunity to choose our Cas9 line too. Under the *nanos* promoter, this gene drive was inherited in around 90% of offspring, but there were medium fitness costs for female carriers, as some offspring was not viable due to the above-mentioned maternal deposition of gene drive components (54). For this study, we thus chose to use a different Cas9 line, namely one that was shown to be highly efficient in males and which showed a lower fitness cost in females (37). This Cas9 is expressed by the *CG4415* promoter (which normally promotes the *gNacα* gene, which is a ribosome associated protein that is predominantly expressed in the male germline (69)), combined with a *nanos* 3’UTR. Both the gene drive and Cas9 constructs are contained within white-eye *w*^*1118*^ lines of *D. melanogaster* called AHDr352v2 and SNc9XnGr, respectively.

### D. Experimental crosses

We follow a standard gene drive crossing scheme with an extra step to introduce genetic variation into the cross, similar to Champer et al. (41) (simplified in Figure 2C). In the first cross, we crossed the two lines containing the split drive elements, the Cas9 construct (with green fluorescence) and the driving construct containing the gRNAs (with red fluorescence), hereafter called the gene drive. The gene drive was contributed by the paternal side of this cross. Secondly, we crossed males heterozygous for both elements of the split drive back to females from the Cas9 line. From those offspring, we selected males carrying the gene drive (red fluorescence) and that were likely homozygous for Cas9 (bright green fluorescence rather than fainter green). Thirdly, we crossed these to females of the genetically diverse DrosEU populations. These DrosEU females were all isolated from separate single crosses in separate vials, making it possible to correct for non-genetic confounding factors such as shared environmental effects or maternal effects (70, 71). From that offspring, we selected five males for the final cross back to *w*^*1118*^ females. These males were likely carrying the gene drive (due to it being driven in their male parent) as well as Cas9 (due to their male parent being homozygous for Cas9). Females were allowed to lay eggs for 7 days. In the offspring, all males had a white-eye phenotype (as the white-eye mutation is X-linked), making it possible to score fluorescence and calculate drive inheritance. Females had wild-type eyes, making fluorescence scoring substantially more difficult. Thus, fluorescence was only scored in male progeny. All offspring, including females, were counted.

Because we were interested in fitness costs due to the gene drive and its activity, besides these gene drive crosses we followed the same crossing scheme with flies that carried Cas9, but not the drive. This allowed us to distinguish any drive-related fitness costs from any DrosEU-*w*^*1118*^ outbreeding fitness costs.

### E. Fly rearing

All crosses except the final cross were performed in bottles with 50 mL of standard medium containing yeast (140 gr/L), sugar (100 gr/L), and agar (20 gr/L), with nipagin (15 mL of 100 gram/L EtOH) and propionic acid (3 mL of 13.39 M) to avoid bacterial or fungal contamination. The final crosses were performed in vials with 6 mL of the above-mentioned medium. Flies were kept at 25°C on a 16:8 hours cycle of light:dark. Handling of flies happened under CO_2_ sedation. Experiments were done in compliance with Dutch genetically modified insect regulations.

### F. GLMM and data analysis

For data visualisation and data analysis, we used R version 4.4.0 (72). For the GLMM variance partitioning as well as data simulation from the fitted model, we used the package glmmTMB (version 1.1.10) (73) in combination with the DHARMa package (version (0.4.7) (74) to test for uniformity of distribution. For details and code, see the Github repository (https://github.com/NickyFaber/HomingDrive_Heritability).

### G. Mutations in *RpL35A* screening

To screen for mutations in *RpL35A* at the gRNA target sites, DNA was isolated from individual flies that did not inherit the gene drive (that is, did not show red fluorescence). Using a high salt protocol, flies were ground in 100 ul buffer (0.1 M Tris HCl pH 8, 1 mM EDTA, 0.25 M NaCl, 1 μg/ul Proteinase K), and incubated overnight at 55 °C. Proteinase K was inactivated by heating for two minutes at 95 °C, and tubes were centrifuged for three minutes at 15.000 rpm to collect debris at the bottom. Supernatant containing the DNA was transferred to a clean tube and diluted 10x. Primers were designed with Primer3 (75) and tested both on *w*^*1118*^ flies and on several wildtype DrosEU flies to check that they worked equally efficiently for both. Primer sequences are 5’GGACGCCTCTTCGCCAAG’3 for the forward primer and 5’GCAAGAAGAAAGAAGGCATGT’3 for the reverse primer. PCR was done using a standard GoTaq polymerase PCR protocol and put on a 1% agarose gel with ethidium bromide. Sanger sequencing was performed by Eurofins to determine heterozygosity. PCR product of identified heterozygous flies was sequenced with Nanopore sequencing technology by Plasmidsaurus.

Sequences were mapped to the *D. melanogaster* reference genome using Minimap2 (76), after which Freebayes was used to call variants (77), Whatshap was used for phasing the two haplotypes (78), and Bcftools was used to create consensus haplotypes (79). A multiple-sequence alignment of the resulting haplotypes was made using the Muscle algorithm in R (version 4.4.0) with the ‘msa’ package (version 1.38.0) (80). All programs were ran with default parameters, but for data and code, see Github repository (https://github.com/NickyFaber/HomingDrive_Heritability).

To check for standing genetic variation at the *RpL35A* gRNA target loci in the DrosEU populations, we used the available DrosEU vcf file containing variant loci and allele frequencies per population (59).

## Results

In total, our experiment consisted of 1343 crosses, 656 of which were gene drive males crosses with *w*^*1118*^ females, and 687 of which were Cas9-only males crossed with *w*^*1118*^ females as control. Fluorescence was scored in all male offspring of the gene drive crosses (female offspring had red wild-type eyes that makes fluorescence scoring difficult, see Figure 2C), and for both the gene drive and control crosses the number of offspring was counted. Finally, 63 offspring from the 30 gene drive crosses with the lowest gene drive inheritance rates were sequenced to identify mutations that are potential simple resistance alleles.

### A. Gene drive inheritance rate

The first high-level phenotype that reflects gene drive efficiency is gene drive inheritance. Plotting the gene drive inheritance rate in male heterozygotes, the most visually striking result is how similarly the gene drive performs across all groups (Figure 3 for gene drive inheritance and Figure S1 for Cas9 inheritance). The absence of a pattern is also clear when plotting the data only by population (Figure S2) or isofemale line (Figure S3). Because there is a significant amount of spread in the number of offspring produced in each cross (see next section), we confirmed that gene drive inheritance is not meaningfully impacted by this due to, for example, larval crowding effects (Figure S4 and Figure S5).

**Figure 3.**
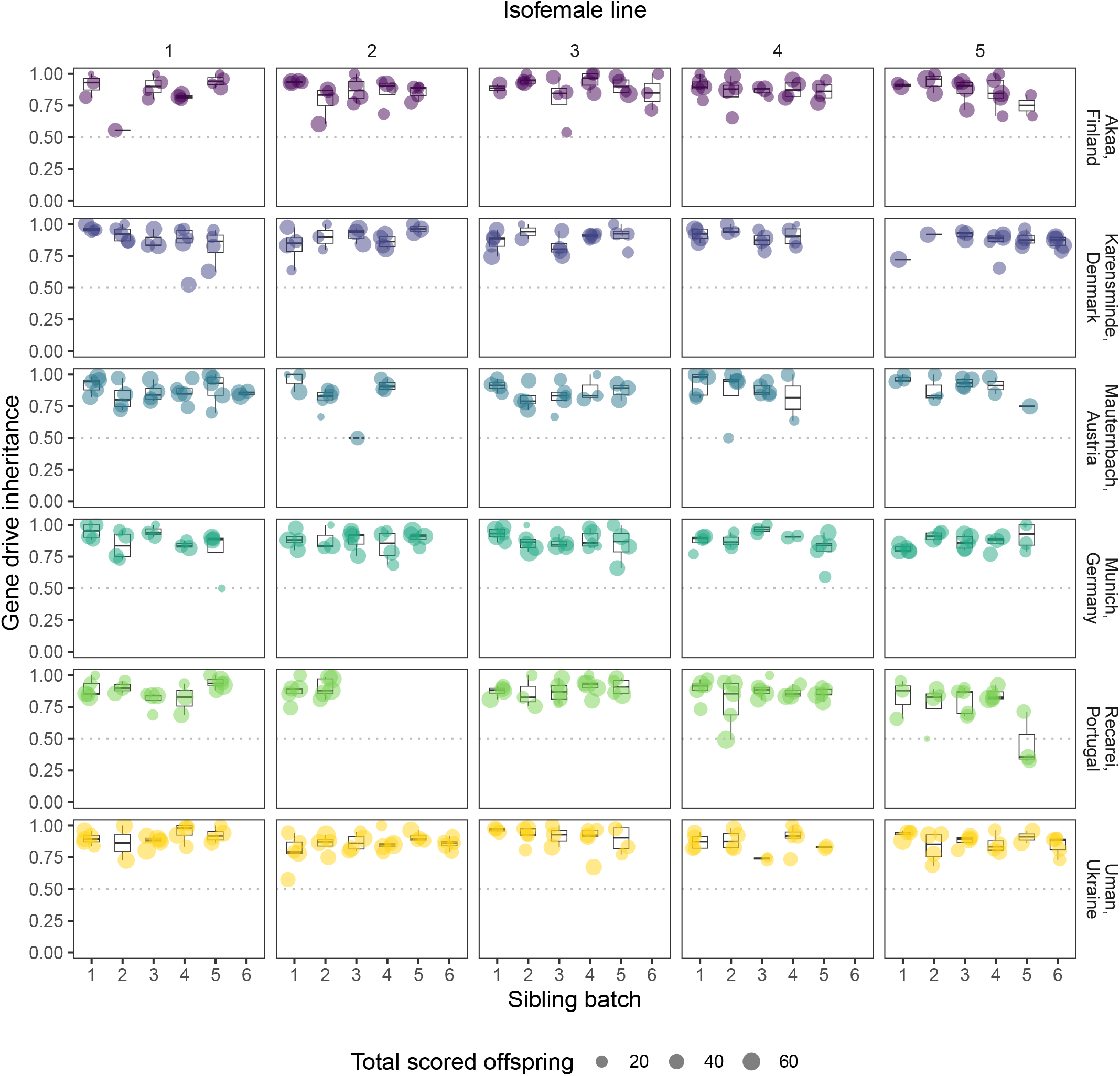
Gene drive inheritance rates plotted per each hierarchical level of the experiment. A dashed line at 0.5 indicates Mendelian inheritance. The size of each dot indicates the number of male offspring in that cross. In total, out of 656 crosses that were made, 579 crosses produced at least one male offspring that could be scored for fluorescence. For the same plot grouped only by higher levels in our experiment (population and isofemale line), see Figure S2 and Figure S3.

Fitting a GLMM containing the three experimental hierarchical levels (population, isofemale line, and sibling batch) as nested random effects, we find that there are significant problems with uniformity of distribution (Asymptotic one-sample Kolmogorov-Smirnov test, p = 0.0027), dispersion (DHARMa nonparametric dispersion test, p = 0.04), and outliers (DHARMa bootstrapped outlier test, p = 0.016). This means that our data significantly diverges from a true binomial distribution due to unaccounted causes of variance. We visualize this in Figure S6, where we show the distribution of our observed data *versus* 100 simulations of a true binomial distribution from the fitted model. To account for the unknown causes of variance in our experiment, we add an observation-level random effect to the model.

After performing model selection by sequentially removing the hierarchical level that explains the least amount of variance, we conclude that the best fitting model keeps only the lowest level (the observation-level random effect, cross). This means that none of the hierarchical levels in our experiment significantly contributes to explaining the variance observed in gene drive inheritance rates. Neither sibling batch (*χ*^2^(1) = 2.23, *p* = 0.14), isofemale line (*χ*^2^(1) = 0.95, *p* = 0.33), or population (*χ*^2^(1) = 0.00, *p* = 1.00) are kept as random effects, nor who (out of four people) scored the flies as a fixed effect (*χ*^2^(3) = 3.50, *p* = 0.32)

(Table S1). From the model containing all four experimental levels, the total amount of variance that can be explained by each level is shown in Figure 4, with effect ranges per group shown in Figure S7. Using the heritability formula specified in Equation 1, we calculate that the heritability of this gene drive’s efficiency (that is, inheritance rate) in the DrosEU lines is 0.31%, though this value is likely not significantly different from 0% as the best fitting model retains neither of the two experimental levels representing genetic effects.

**Figure 4.**
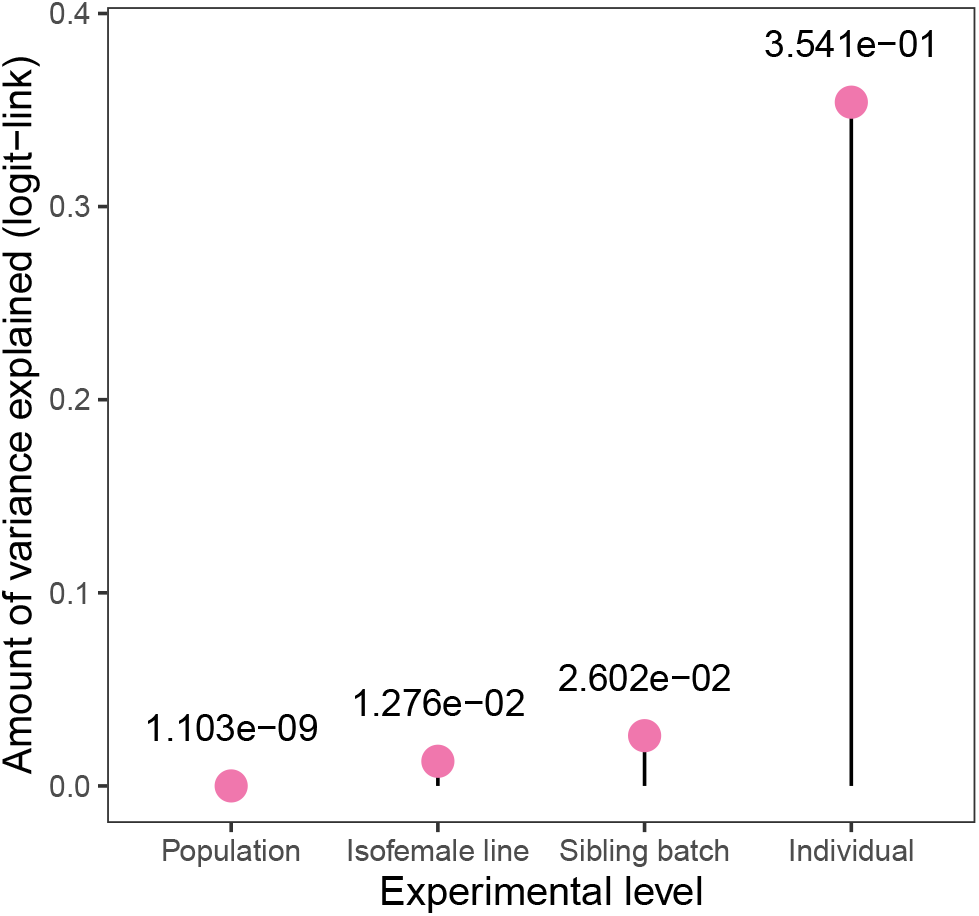
Variance in gene drive inheritance rate partitioned to each hierarchical level of our experiment. These values are logit-linked.

### B. Number of offspring

The second factor determining gene drive efficiency is fitness of gene drive carriers, which we measure here as the total number of adult offspring. Because *RpL35A* is a haplolethal gene, any offspring inheriting a non-functional allele due to indels will likely not be viable (54). Furthermore, just the presence of active Cas9 could lead to a fitness costs, likely due to off-target cutting (81). When plotting the overall number of offspring from gene drive crosses and control crosses (which only carried Cas9, and not the gene drive itself), we see that the average number of offspring is similar between gene drive crosses (mean = 68.15, n = 656) and control crosses (mean = 68.89, n = 687). However, the distribution appears wider in both directions for gene drive crosses, indicative of the gene drive causing some fitness effects (Figure 5). When plotting the data per population (Figure S8), per isofemale line (Figure S9), or per sibling batch (Figure S10), subtle differences in the number of offspring between isofemale lines are visible, but there is a large amount of overall variance.

**Figure 5.**
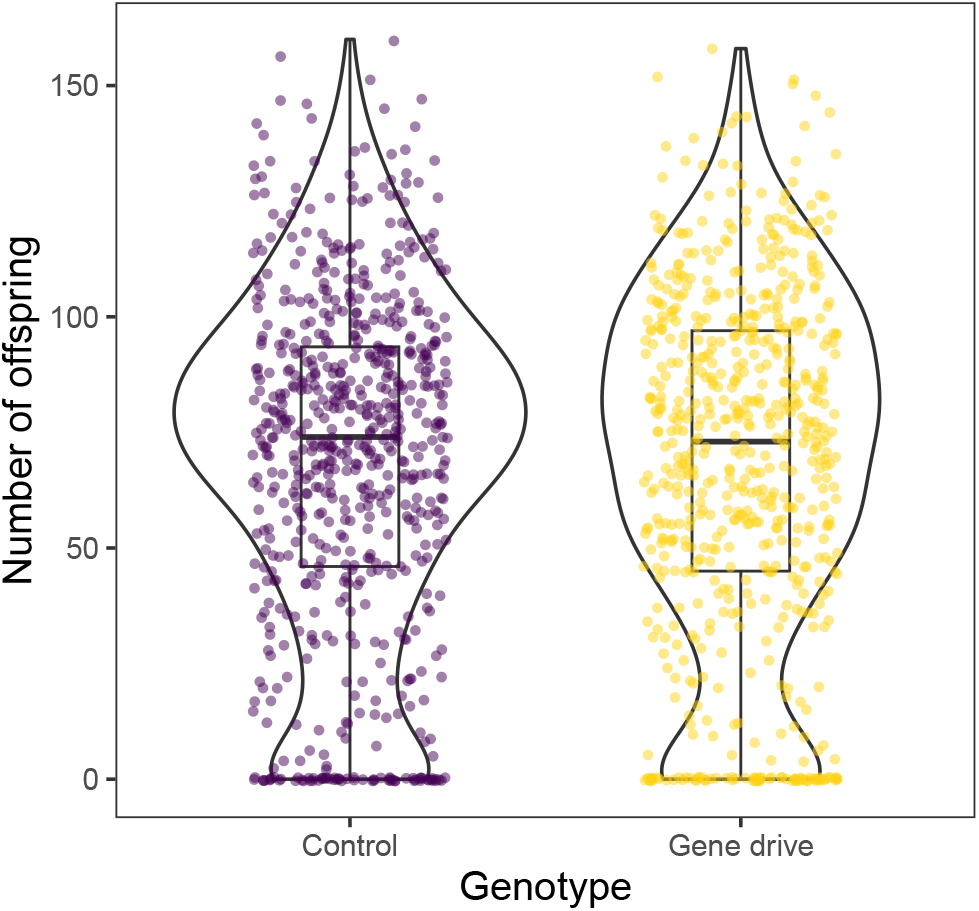
Number of offspring from gene drive and control crosses. Violin plots are drawn with default “nrd0” smoothing and are proportionally sized to the number of replicates. The data is additionally plotted per population in Figure S8, per isofemale line in Figure S9, and per sibling batch in Figure S10.

Fitting a GLMM that corrects for zero-inflation (as some crosses produced no offspring), we find that a Gaussian distribution with a zero-inflation model fits our data the best (Figure S11). The full model contains the three experimental hierarchical levels (population, isofemale line, and sibling batch) as nested random effects, and genotype (gene drive or control) as a fixed effect, as well as interactions between genotype and population or line. This interaction indicated any gene drive effect that is specific to a population or line. The difference in heterogeneity of variance between control and gene drive crosses is significant (Levene’s Test for Homogeneity of Variance, p < 0.032) (Figure 5). Otherwise, the distribution of the data is normal in terms of uniformity (Asymptotic one-sample Kolmogorov-Smirnov test, p = 0.26) and dispersion (DHARMa nonparametric dispersion test, p = 0.8), although there are fewer outliers than expected, probably due to the zero-inflation model (DHARMa bootstrapped outlier test, p < 0.001). We do model selection by sequentially removing the effect that explains the least amount of variance. The best fitting model contained only isofemale line (*χ*^2^(1) = 45.60, *p <* 0.001), but not population (*χ*^2^(1) = 0.57, *p* = 0.45), or any of the interactions between population and isofemale line (see Table S2 and Figure S12). Isofemale line is expected to be a significant factor, as setting up isofemale lines creates significant genetic bottlenecks (60). Overall, we conclude that there is no meaningful interaction between genetic background and the fitness effect of the gene drive (measured as the total number of offspring). However, due to the large amount of total variance we observe, perhaps a more targetted experimental set-up than the one used in this study is necessary to uncover potential small effects.

### C. Resistance alleles in RpL35A at the gRNA target sites

We investigated the crosses with the lowest gene drive inheritance for potential simple resistance alleles. These alleles could already have been present in the DrosEU populations, or they could have been formed through gene drive activity. Based on population genetic data from the DrosEU consortium, we do not expect any standing variation as there is not a single variant observed at either of the two gRNA sites in any population across Europe (Figure S13A and B). Furthermore, plotting the frequency of variants in the whole *RpL35A* gene, we see that most variants are at low frequency in most populations (Figure S13C).

Therefore, variants in the homology regions during gene drive homing seemed unlikely, but not an impossible, cause for the occasionally strongly reduced homing efficiencies (82, 83). At the same time, we assumed resistance allele formation through gene drive activity itself to be highly unlikely too due to the conserved and haplolethal nature of the RpL35A target gene, together with the use of two gRNAs (54).

From each of the 30 lowest-drive inheritance crosses, we took up to three males that did not inherit the gene drive (that is, they were not red fluorescent), and amplified and sequenced roughly 400 bp of the *RpL35A* locus (Figure S13A). Surprisingly, out of 63 flies in total, 11 flies were heterozygous at the *RpL35A* locus, whereas the rest was homozygous wildtype (Figure 6). The resistance alleles were never the same between siblings of a single cross, and were also not disproportionally found in any particular population or isofemale line. These 11 resistance alleles can be classified into two types: 1) double deletions at both gRNA sites and 2) partial homology-directed repair from the gene drive’s *RpL35A* rescue element. Even more surprisingly, out of the six haplotypes with double deletions, only three leave *RpL35A* in frame, and the other three haplotypes have a frame-shift in *RpL35A*. Although *RpL35A* is assumed to be a haplolethal gene, these flies were viable despite this frame-shift, because they lived to adulthood before they were frozen at the end of the experiment. Because of this, the fertility of these flies could not be checked and we cannot say whether they are functional resistance alleles (that is, potentially transmitted to the next generation).

**Figure 6.**
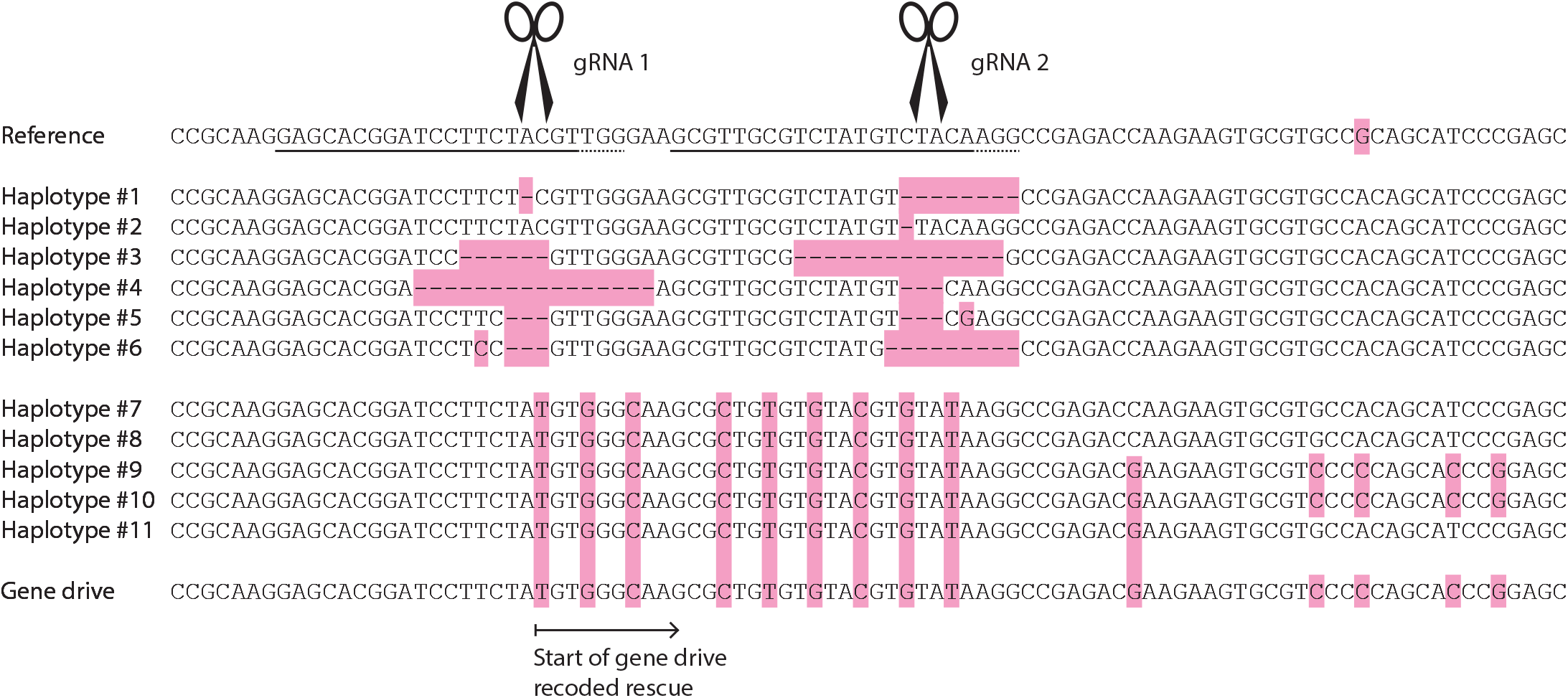
Multiple sequence alignment of the 11 observed RpL35A resistance alleles. On top is the *D. melanogaster* reference genome, where we show the gene drive’s two gRNAs targets (solid underscore indicates the 20 bp gRNA target sequence, dashed underscore indicates the PAM-sequence, and scissors indicate the location where the double-stranded break will occur. Below the reference sequence, we show the 11 resistance allele haplotypes that were found, with pink blocks indicating deletions and mismatches with the reference. At the bottom, we show the gene drive sequence. Note that haplotypes number 9 and 10 contain a slightly larger piece of the recoded sequence than is shown here; for the full alignment, see the GitHub repository (https://github.com/NickyFaber/HomingDrive_Heritability).

Of the five haplotypes that showed partial homology-directed repair from the drive allele (as can be seen by the substitution roughly every third base, which was designed to give drive individuals a functioning *RpL35A* rescue, without it being susceptible to cutting by the gRNAs), each showed varying lengths of the recoded part of the gene drive (Figure S14); see the GitHub repository for the full alignment, https://github.com/NickyFaber/ HomingDrive_Heritability. Of these, we also cannot conclusively say that they are functional resistance alleles, although arguably these are highly likely to be functional *RpL35A* copies as gene drive individuals are fully fertile with the entire recoded sequence. These genotypes are also likely not susceptible to cutting by the gRNAs because they contain several mismatches at the PAM-proximal end, where mismatches are less tolerated (84). These data show that viable resistance alleles are possible in conserved genes such as *RpL35A*. All in all, this is the first demonstration that partial homology-directed repair could facilitate simple resistance allele formation for gene drives containing a recoded rescue element.

## Discussion

As with any biocontrol method, it is crucial to anticipate the evolution of resistance—regardless of its form—and develop appropriate mitigation strategies (44–50). The well-documented and expected form of resistance against gene drives arises from mutations at its gRNA target sites, which are highly likely to occur and necessitate countermeasures (12–15, 22). In contrast, complex forms of resistance remain understudied and their potential impact on gene drive efficacy is not yet well understood (22, 41). In this study, we investigate the potential for complex resistance to emerge against a CRISPR-based homing gene drive. We do so by testing a modification rescue drive in diverse genetic backgrounds in a nested full-sibling design to estimate heritability, while accounting for non-genetic confounding factors (70). We assessed two key indicators of gene drive efficiency: the inheritance rate of a homing rescue gene drive and the fitness of its carriers as the total number of offspring. Additionally, we screened for mutations that could indicate the presence of simple resistance alleles.

Our findings reveal an exceptionally low heritability, 0.31%, for the inheritance rate of the homing rescue gene drive in male carriers—far lower than typical heritable traits in *D. melanogaster* (85). Several factors may be behind this low heritability. Firstly, although intuitively it makes sense to think of gene drive efficiency as having a genetic component, the gene drive may just be a mostly self-contained construct that relies on gene expression machinery. Indeed, different Cas9 promoters show more significant variation in gene drive inheritance rates than we observed here (37, 86). Secondly, perhaps genetic effects are so small that our experimental set-up is not sensitive enough to pick them up. As complex cellular mechanisms underlie gene drive efficiency (Cas9 expression, Cas9 protein activity, DNA accessibility, and DNA repair mechanisms), this seems possible (6, 13). Thirdly, these mechanisms may be highly conserved, thus limiting variation. However, there is evidence for genetically determined variation in somatic DNA repair, indicating that this might be possible in the germline too (87). Finally, potential trade-offs between DNA repair pathways and fitness could constrain observable differences. For example, a shift from homology-directed repair to end-joining repair in the germline could lead to a direct fitness cost because of the necessary double-stranded break repairs during crossing-over, thus still keeping overall drive inheritance similar by removing wild-type alleles instead of turning them into drive alleles.

The second part of gene drive efficiency we assessed was the total number of offspring. End-joining could lead to lethal genotypes in the offspring, an effect that would be visible in a reduction in offspring numbers. Although we saw a noticeable, though small, shift in the distribution, this effect was likely confounded by larval crowding effects and thus hard to statistically analyse. Further, fecundity is a highly variable trait in *D. melanogaster* (88), so observing a reduction in the number of offspring is harder to measure than the occurrence of a visible phenotype such as fluorescence or a yellow body colour (13).

The low observed heritability, despite substantial observed variation, suggests that selection for complex resistance against this gene drive in genetically diverse populations will be inefficient (89). This is beneficial for gene drive efficacy, as it implies this gene drive will likely remain relatively stable over generations. However, with a large enough selection pressure, for example, if this gene drive were converted to a suppression drive (32), and a large enough population, complex resistance could still be selected for. Before field trials are initiated, it is essential to screen gene drives specifically in the genetic background of the target population to better understand the potential for complex resistance across different constructs, populations, and species. While field trials could reveal the effects of complex resistance, they also carry the risk of irreversibly altering the genetic makeup of wild populations. Although our findings suggest that selection for complex resistance is likely inefficient, resistance mechanisms such as Cas9 inhibition or shifts in DNA repair pathways would impact all gene drives relying on these processes. Unlike simple resistance, these forms of resistance cannot always be mitigated by modifying gRNAs or designing a new gene drive (28). This consideration should be incorporated into risk assessments and public engagement efforts prior to field deployment. Regular sampling of heterozygous gene drive individuals from genetically diverse populations and testing their gene drive efficiency in the lab could provide ongoing monitoring of drive functioning.

Although our primary focus was to investigate complex resistance, we also identified multiple resistance alleles at the gRNA target sites in *RpL35A*. These resistance alleles were somewhat unexpected given the conserved nature of this gene, which codes for a ribosomal subunit that was believed to be haplolethal, though mosaic resistance was seen previously (54) (see previous modelling of partial repair (27)). Based on the mutations we found in *RpL35A*, some of which were out of frame, it seems likely that this gene is not completely haploinsufficient, perhaps at least allowing some mutations in some genetic backgrounds to be viable (66). Although we cannot determine whether these mutations confer functional simple resistance, it could be that they are at least recessively lethal. However, we believe that the resistance alleles copying some of the recoded part of the gene drive (replacing the original wild-type sequence) may represent functional resistance as gene drive individuals are also fully viable and fertile with the full recoded sequence.

These findings raise a key question: will gene drive resistance in natural populations primarily arise as simple mutations, complex mechanisms, or perhaps not at all? Our data suggest that simple resistance will be the more immediate concern, and that homing rescue gene drives have an additional mechanism through which they are vulnerable to the formation of these alleles. It is not immediately clear how these alleles can be mitigated. One option is through multiplexing with more than two gRNAs, as modelled by Champer et al. (27). Another option is to target conserved genes with less functional redundancy than RpL35A such as *doublesex* (31). In any case, if this gene drive were converted to a suppression drive, this type of two-target suppression drive is more tolerant of simple resistance alleles at the homing site, as the genetic load is mostly imposed at the distant target (32). Adjusting the 3’ UTR to a version from another species (with several mutations to break up the large tract of identical DNA) could both improve drive efficiency and reduce the chance of partial homology-directed repair (though it is unclear to what extent the 3’ UTR is involved in the partial homing process), though it might also reduce the chance of successful rescue (58). Using genes with smaller 3’ UTRs may also reduce functional resistance rates.

Future research could explore several avenues. Related to the evolution of complex resistance, our conclusion comes with the caveats that our study examined only a single gene drive construct, only in males, and only exclusively in European *D. melanogaster* populations. Regarding the gene drive and promoter used in this study, a less efficient gene drive or one that is not a haplolethal rescue construct may reveal greater variation across genetic backgrounds, although these inefficient constructs are less attractive for use in practice (37, 41). Regarding the amount of genetic diversity tested here, future studies could incorporate a broader range of genetically diverse lines, such as the global diversity lines (90), to better assess the evolution of complex resistance. However, measures of overall nucleotide diversity are only slightly higher in African and North-American populations of *D. melanogaster* compared to European populations (91). Furthermore, a few studies have examined the impact of genetic diversity on gene drive efficiency, but none of them have reported major effects (38–43). Champer et al. (41) conducted a genome-wide association study (GWAS) using the Drosophila Genetic Reference Panel and reported highly heritable effects on several aspects of gene drive efficiency (particularly embryo resistance allele formation rate, and potentially cutting and homing rate). Notably, however, their analysis did not explicitly account for maternal effects or cross-level effects, not ruling out that most of that heritability was confounded by these factors (70). Interestingly, however, embryo resistance rate effects appeared consistent in some lines when using a different gene drive (41). Embryo resistance levels also tended to have intermediate values (on the binomial scale from zero to one), potentially allowing for genetic variation to induce larger absolute effect sizes on this performance aspect. Taking together all this evidence, we think that future studies should assess a range of gene drive constructs and effects, in both males and females, rather than testing more diverse populations of *D. melanogaster*.

Unrelated to the evolution of resistance, we think that our data points us in a direction for improving homing gene drives. Gene drive activity is usually assumed to occur independently for each offspring during meiosis, when homologous chromosomes are paired (13). In our hierarchical experiment, however, a large amount of variance could be attributed to sibling batch and cross-level effects. Therefore, we propose that there are developmental processes relevant for gene drive efficiency that require consideration. This hypothesis aligns with previous indications that gene drives may already be active in pre-meiotic germline stages, potentially skewing inheritance rates in either direction which results in this kind of variation (6, 13, 36, 92, 93). Limiting the timing of a gene drive to the most optimal window for homology-directed repair, while considering the entire developmental cycle of the species, could probably yield efficiency improvements as well as reduce the amount of variance in efficiency. Regardless of the underlying cause, future gene drive studies should consider correcting for sibling batch (maternal and environmental effects) and cross-level effects in their experimental designs, or investigating them explicitly to better understand their potential impact on drive efficiency.

Overall, this study highlights that homing gene drives are likely still their own worst enemy, as mutations and partial homing events potentially indicative of functional simple resistance were found even in a highly conserved target gene. Furthermore, we demonstrate that complex resistance is unlikely to be a challenge for this specific gene drive, at least in males for drive efficiency. Whether this finding extends to other scenarios remains to be determined. As there are currently some homing gene drives under consideration for field trials, we suggest that these could be opportune moments to look for complex resistance in a field setting. This screening is important, as complex resistance potentially still poses a long-term threat with large consequences for future gene drive applications in that population. Finally, we demonstrate that to test the heritability of complex resistance, an experimental design is necessary that accounts for the many potential confounding factors, the causes of which are interesting in and of itself. Our work contributes to testing the robustness of gene drive technology in real-world settings, bringing us one step closer to harnessing their full potential for genetic biocontrol.

## ACKNOWLEDGEMENTS

We are grateful to the Dutch Graduate School for Production Ecology & Resource Conservation (PE&RC) for financially supporting this work. A massive thank you to Gabriella Bukovinszkine Kiss, Jordy Litjens, Francisca Reyes Marquez, Fleur Keizer, Jimmie Zeelenberg, and Reggie King for help with the experimental work, and thank you to José van de Belt for patience with our fluorescence microscope siege. Thank you to the DrosEU consortium for collecting, maintaining, and sharing the European *Drosophila melanogaster* isofemale lines, as well as Nicolas Rode specifically for helpful discussions and advice. Thank you to Gerrit Gort for help with the statistical analysis, and thank you to the staff at Unifarm for logistical support.

## DATA AVAILABILITY

Data is available on GitHub (https://github.com/NickyFaber/HomingDrive_Heritability).

## CODE AVAILABILITY

Code is available on GitHub (https://github.com/NickyFaber/HomingDrive_Heritability).

## AUTHOR CONTRIBUTIONS

N.R.F., B.A.P., B.J.Z., and J.H. conceptualised the study, and all authors designed the experiments together. Experimental work, analysis, and writing were carried out by N.R.F., with support from J.C., B.A.P., B.J.Z., and J.H. All authors read and reviewed the manuscript.

## Supplementary material

**Figure S1.**
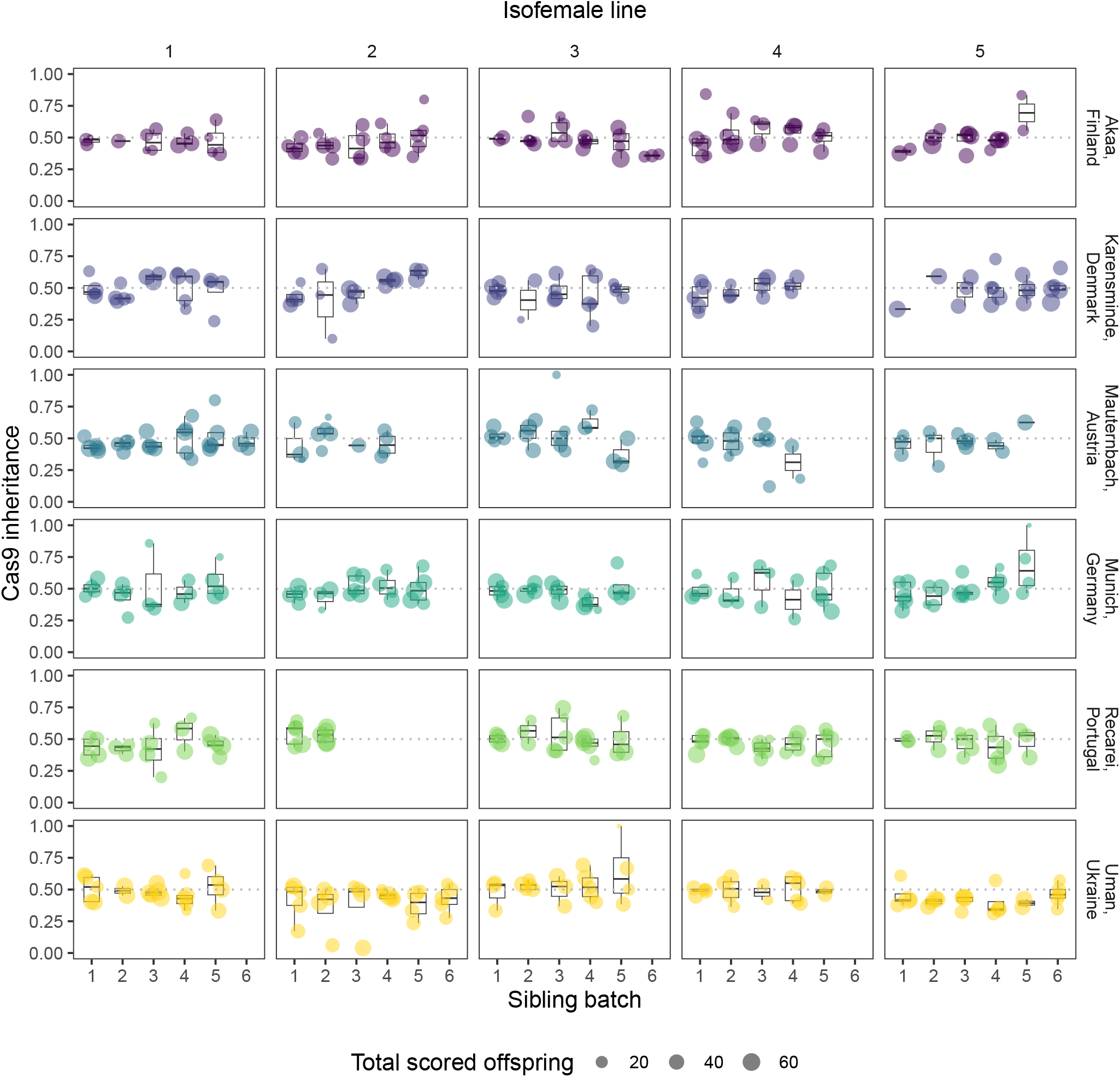
Cas9 inheritance rates plotted per each hierarchical level of the experiment. A dashed line is set at 0.5 to indicate a Mendelian inheritance rate. The size of each dot indicates the number of male offspring in that cross. In total, out of 656 crosses that were made, 579 crosses produced at least one male offspring that could be scored on fluorescence.

**Figure S2.**
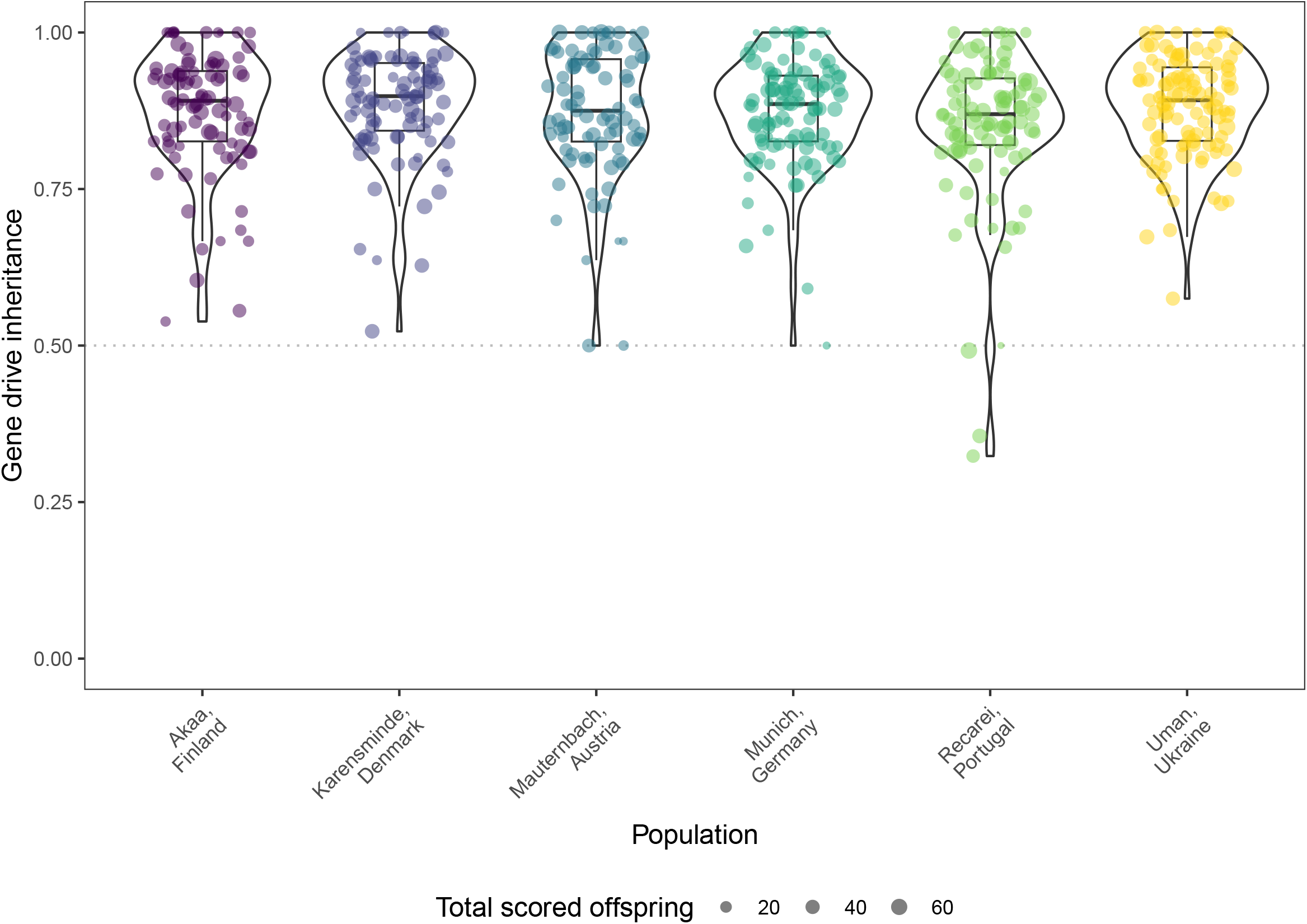
Gene drive inheritance rates plotted per population. A dashed line is set at 0.5 to indicate a Mendelian inheritance rate. The size of each dot indicates the number of male offspring in that cross. In total, out of 656 crosses that were made, 579 crosses produced at least one male offspring that could be scored on fluorescence. Violin plots are drawn with default “nrd0” smoothing and are proportionally sized to the number of replicates. For the same plot grouped by all levels in our experiment, see Figure 3.

**Figure S3.**
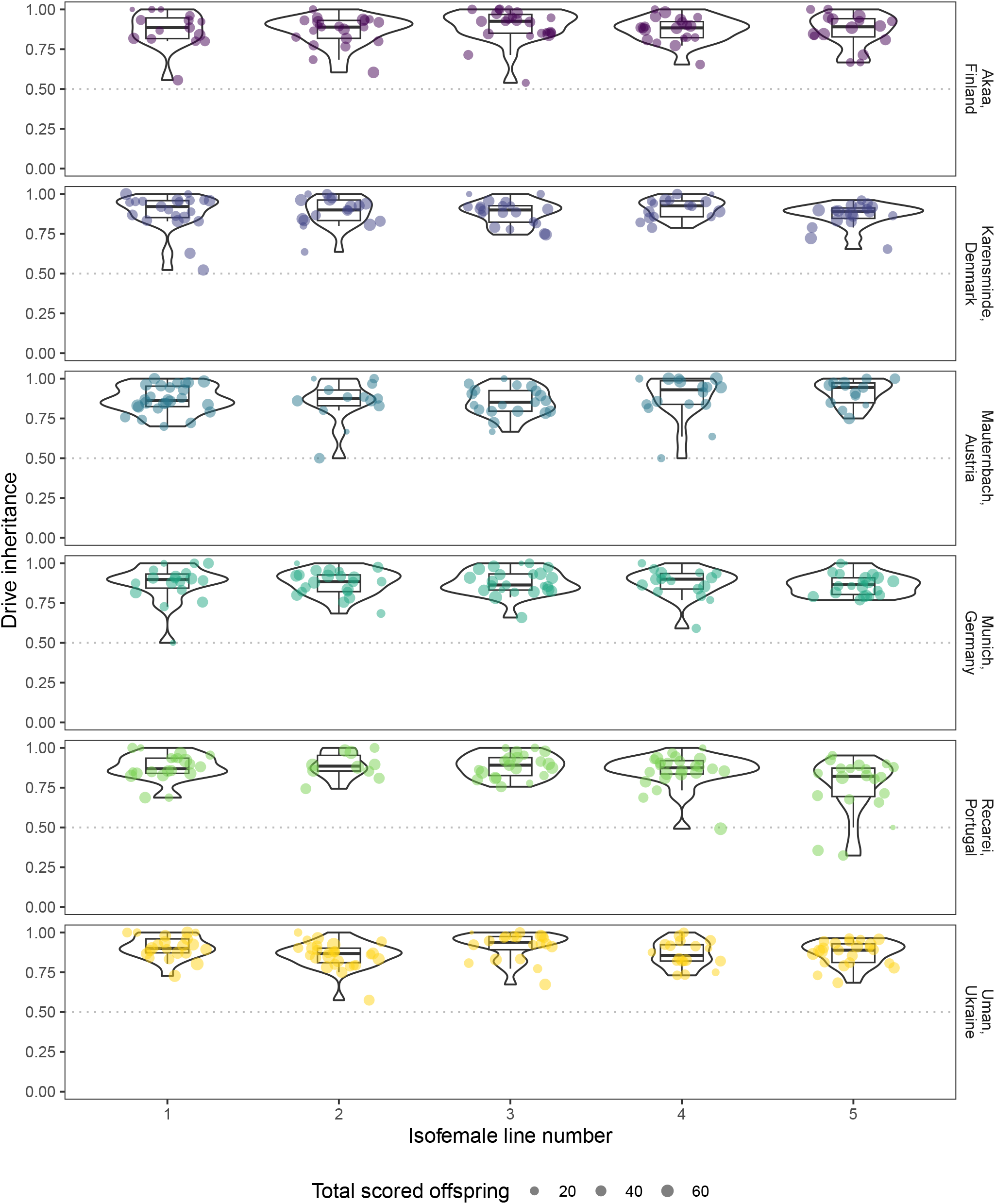
Gene drive inheritance rates plotted per population and isofemale line. A dashed line is set at 0.5 to indicate a Mendelian inheritance rate. The size of each dot indicates the number of male offspring in that cross. In total, out of 656 crosses that were made, 579 crosses produced at least one male offspring that could be scored on fluorescence. Violin plots are drawn with default “nrd0” smoothing and are proportionally sized to the number of replicates. For the same plot grouped by all levels in our experiment, see Figure 3.

**Figure S4.**
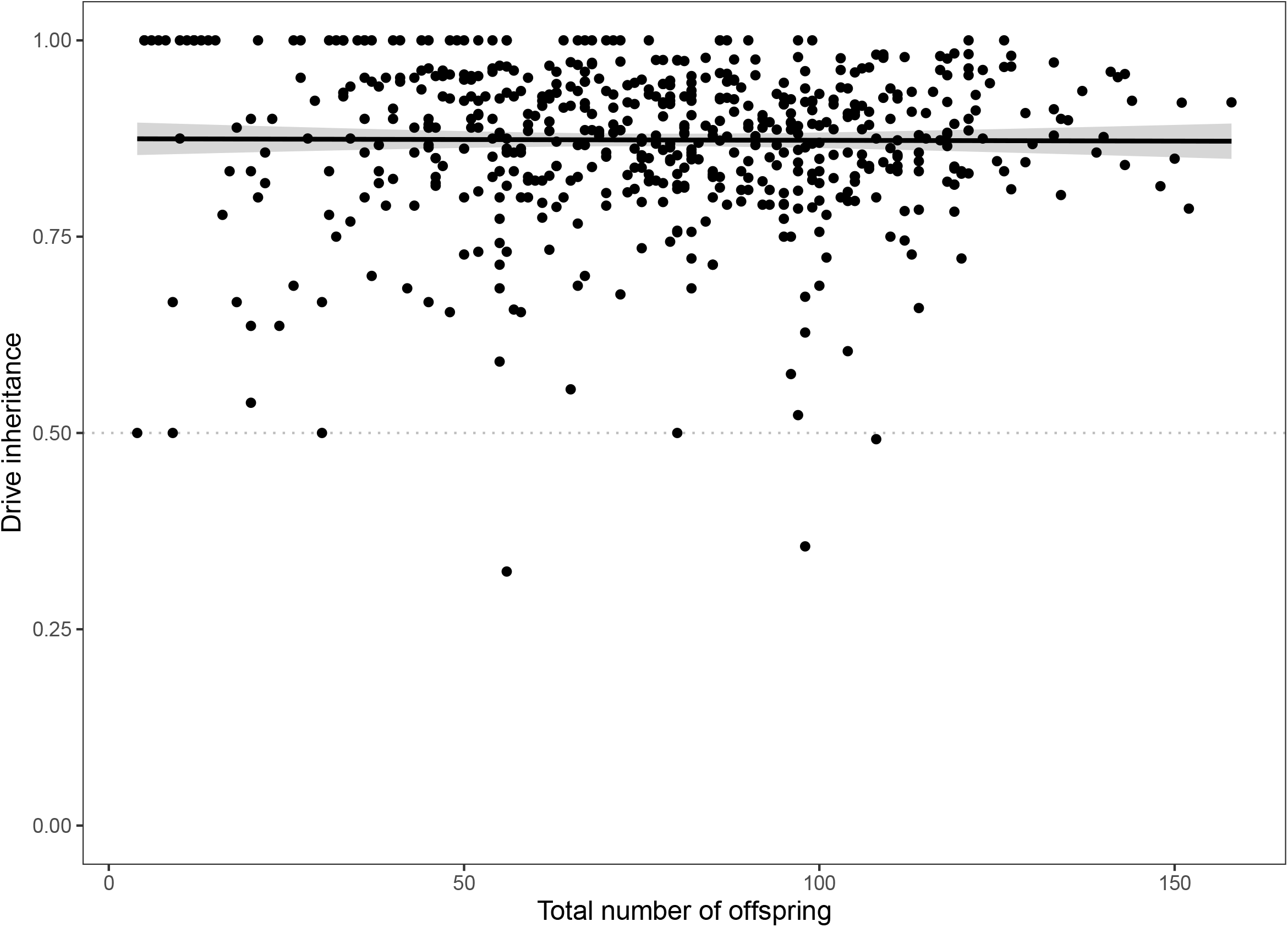
Relationship between gene drive inheritance rate and total number of offspring. Total number of offspring include all male and female offspring, as well as flies that were dead upon collection and were unable to be scored for fluorescence. The line shows the linear regression (using “lm” with formula ‘y x’) and the shaded area shows the 95% confidence interval.

**Figure S5.**
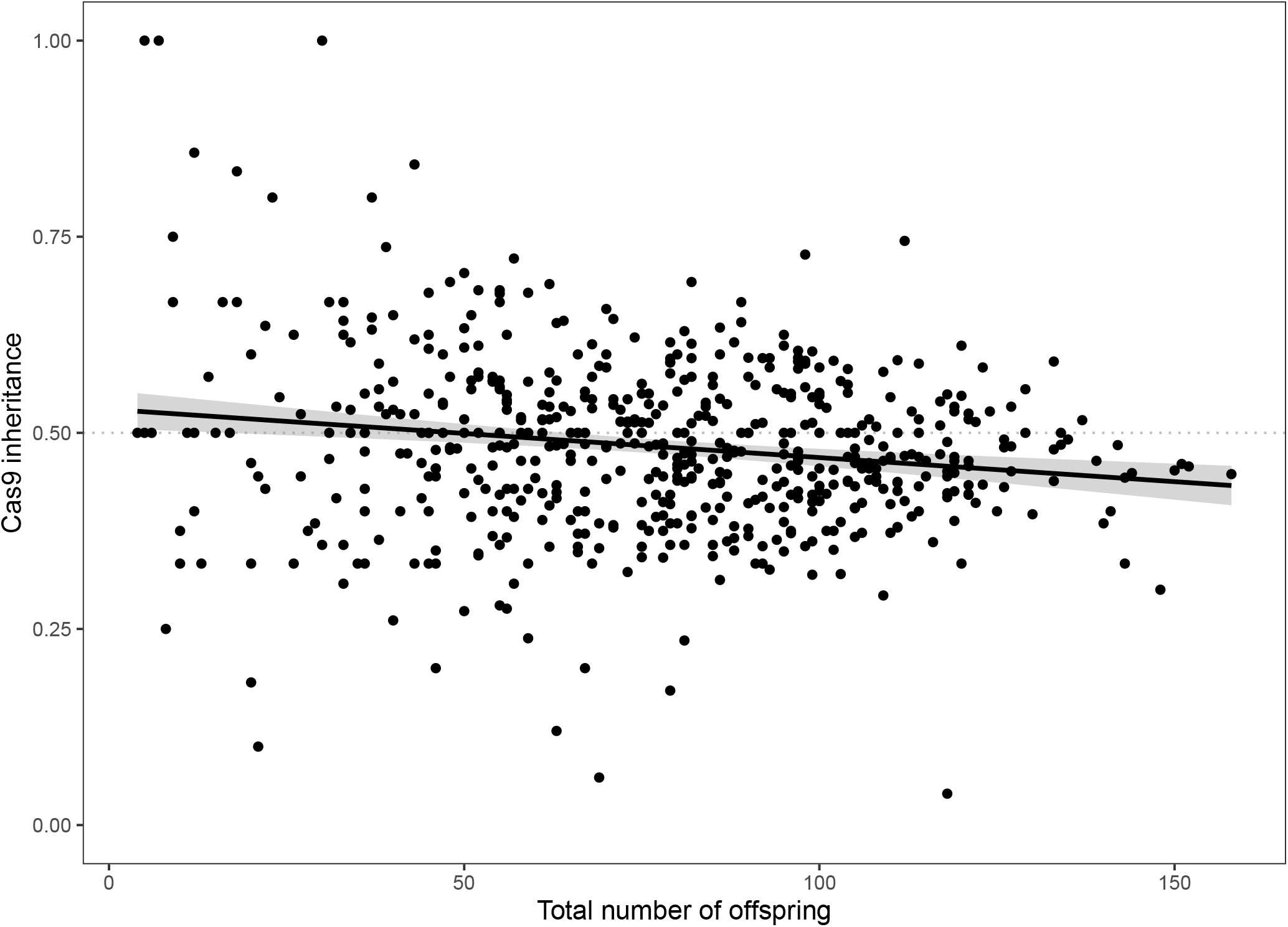
Relationship between Cas9 inheritance rate and total number of offspring. Total number of offspring include all male and female offspring, as well as flies that were dead upon collection and were unable to be scored for fluorescence. The line shows the linear regression (using “lm” with formula ‘y x’) and the shaded area shows the 95% confidence interval.

**Figure S6.**
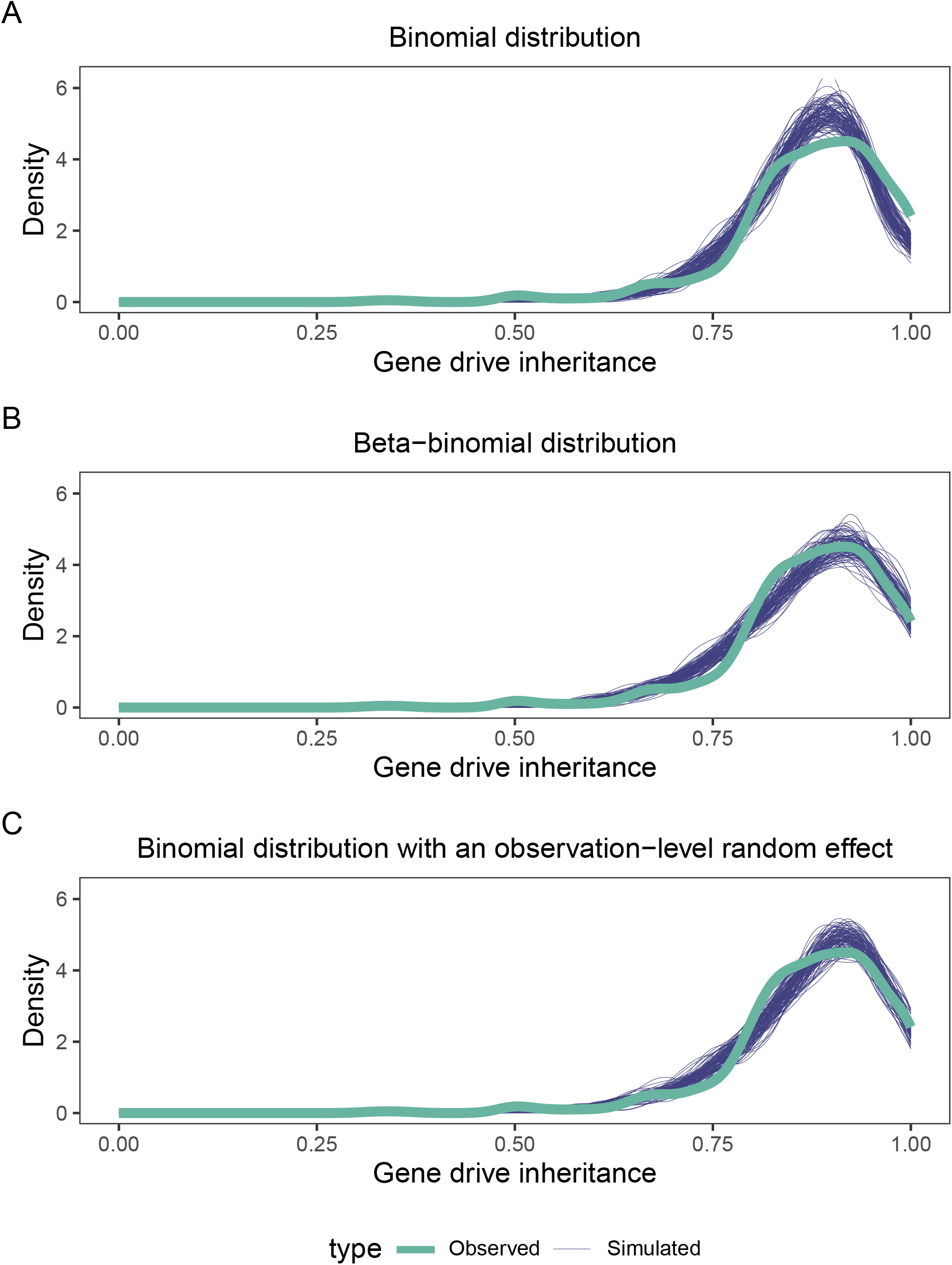
The distribution of gene drive inheritance rates in our data and those from 100 simulations of varying distributions. Simulations were done from a GLMM containing the full hierarchical model with different assumed distributions: **A)** binomial, **B)** beta-binomial, and **C)** binomial with an observation-level random effect.

**Table S1.** Model selection for gene drive inheritance. “OLR” stands for ‘Observation-Level Random effect’.

| Model | Df | AIC | BIC | logLik | deviance | Chisq | Df | Pr(>Chisq) |
| --- | --- | --- | --- | --- | --- | --- | --- | --- |
| OLR_cross | 2 | 2767.8 | 2776.5 | -1381.9 | 2763.8 |  |  |  |
| OLR_batch_cross | 3 | 2767.5 | 2780.6 | -1380.8 | 2761.5 | 2.2340 | 1 | 0.1350 |
| OLR_line_batch_cross | 4 | 2768.6 | 2786.0 | -1380.3 | 2760.6 | 0.9469 | 1 | 0.3305 |
| OLR_population_line_batch_cross | 5 | 2770.6 | 2792.4 | -1380.3 | 2760.6 | 0.0000 | 1 | 1.0000 |
| OLR_full | 8 | 2773.1 | 2808.0 | -1378.5 | 2757.1 | 3.4997 | 3 | 0.3208 |

**Figure S7.**
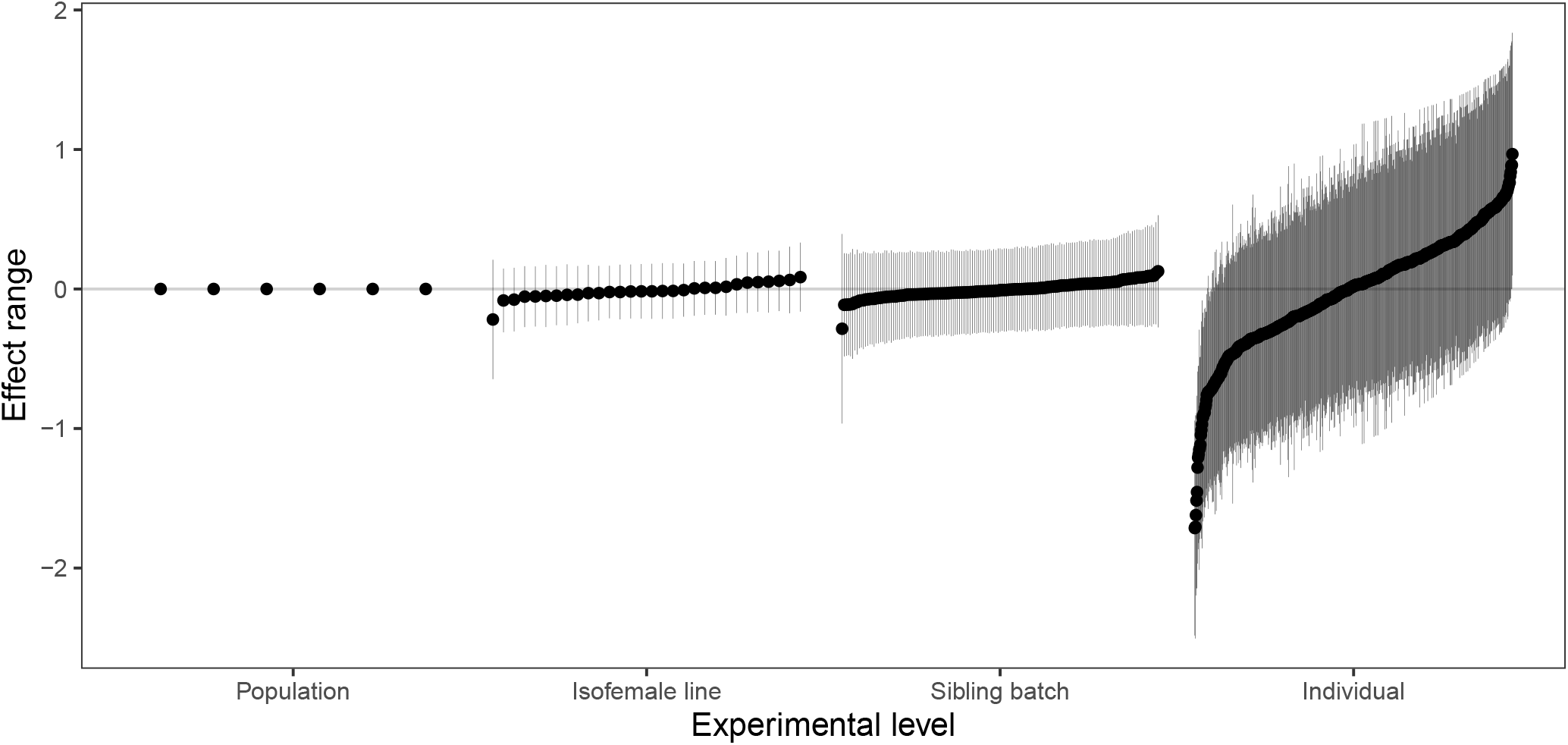
Effect ranges in each group per experimental level. Dots show the estimate of the effect of that group on gene drive inheritance (in log-link) and line ranges show the 95% confidence intervals.

**Figure S8.**
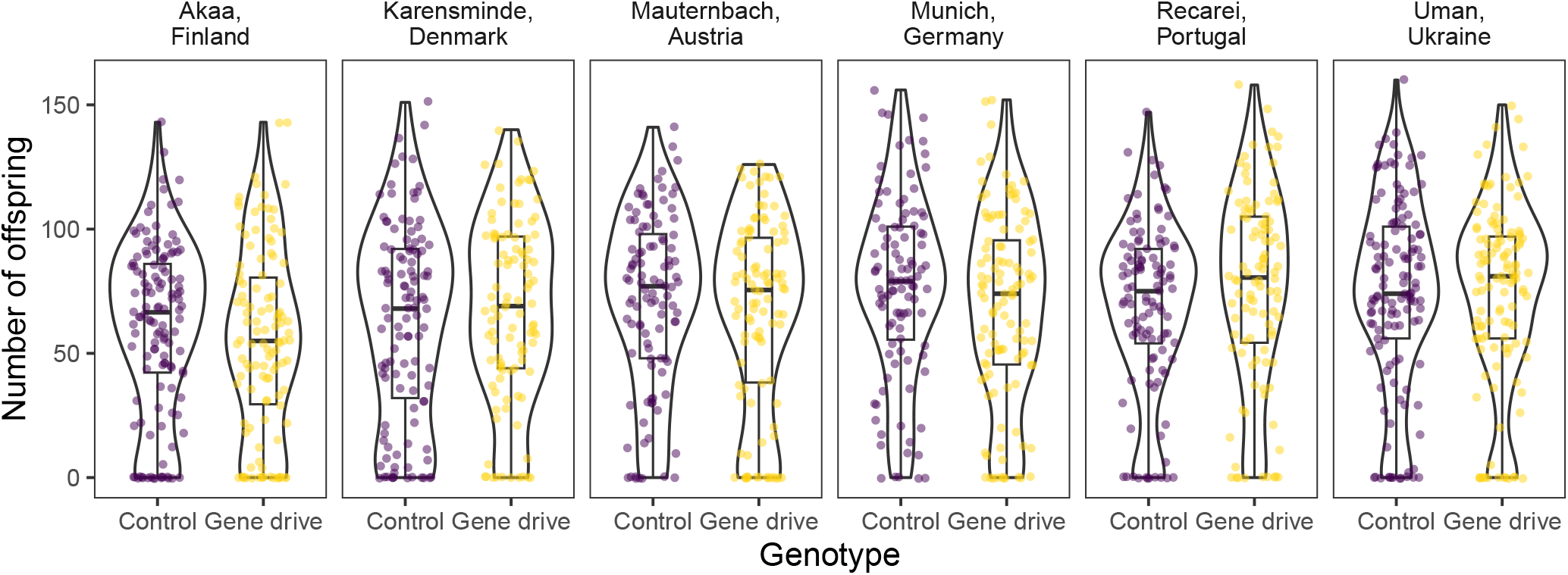
Number of offspring from gene drive and control crosses per population. Violin plots are drawn with default “nrd0” smoothing and are proportionally sized to the number of replicates. The data is plotted per genotype in Figure 5.

**Figure S9.**
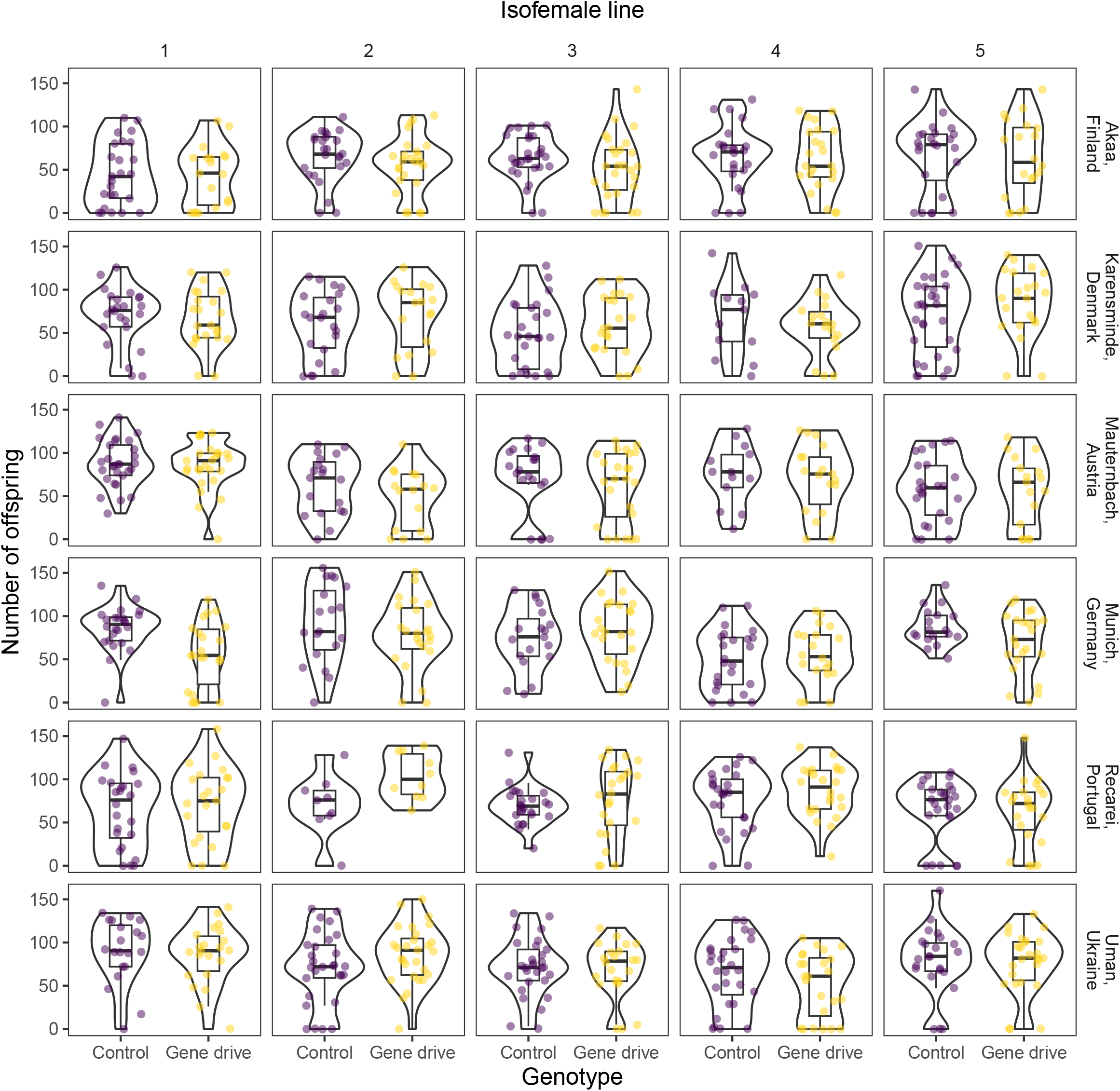
Number of offspring from gene drive and control crosses per population and isofemale line. Violin plots are drawn with default “nrd0” smoothing and are proportionally sized to the number of replicates. The data is plotted per genotype in Figure 5.

**Figure S10.**
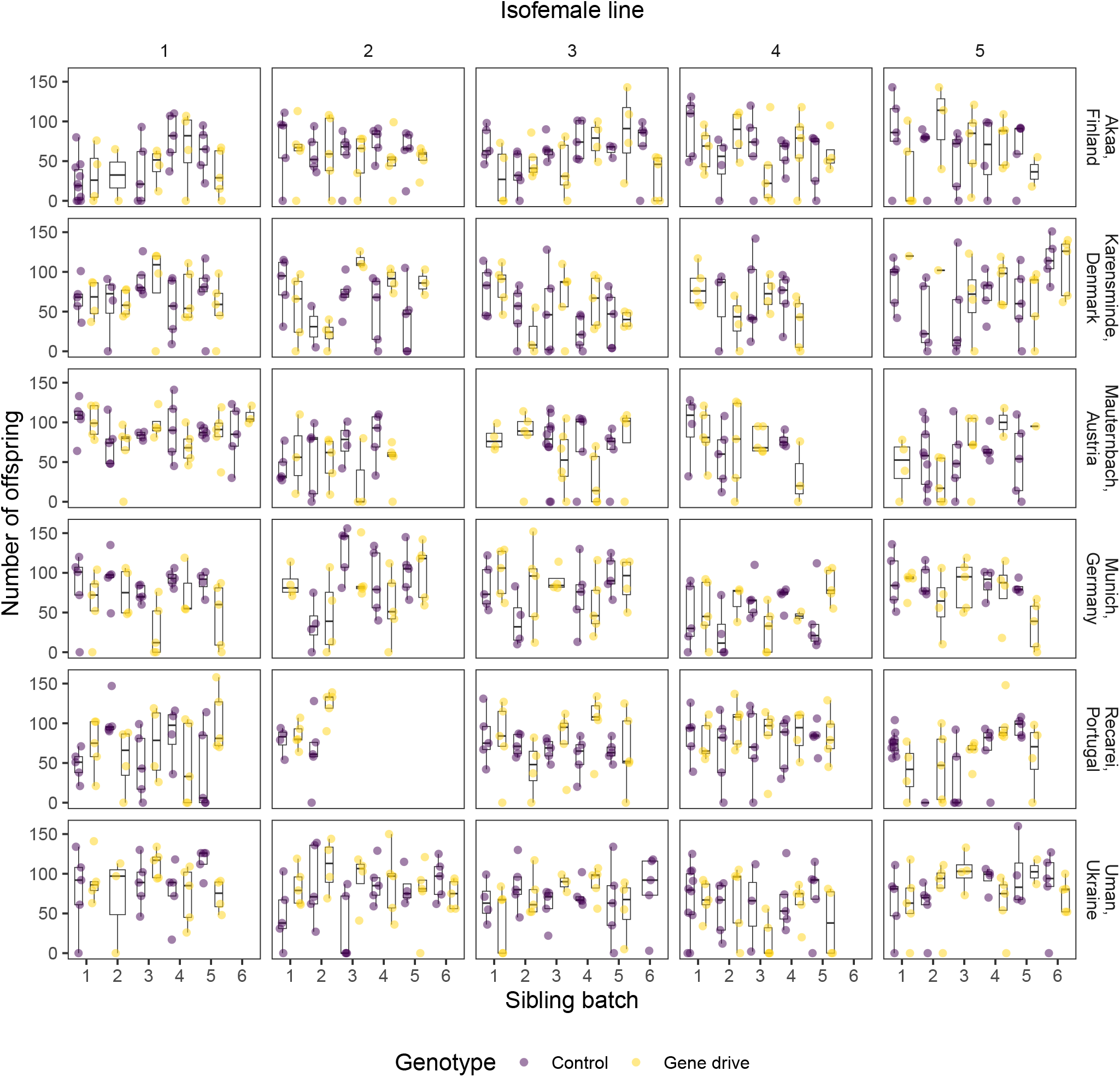
Number of offspring from gene drive and control crosses per population, isofemale line, and sibling batch. Violin plots are drawn with default “nrd0” smoothing and are proportionally sized to the number of replicates. The data is plotted per genotype in Figure 5.

**Figure S11.**
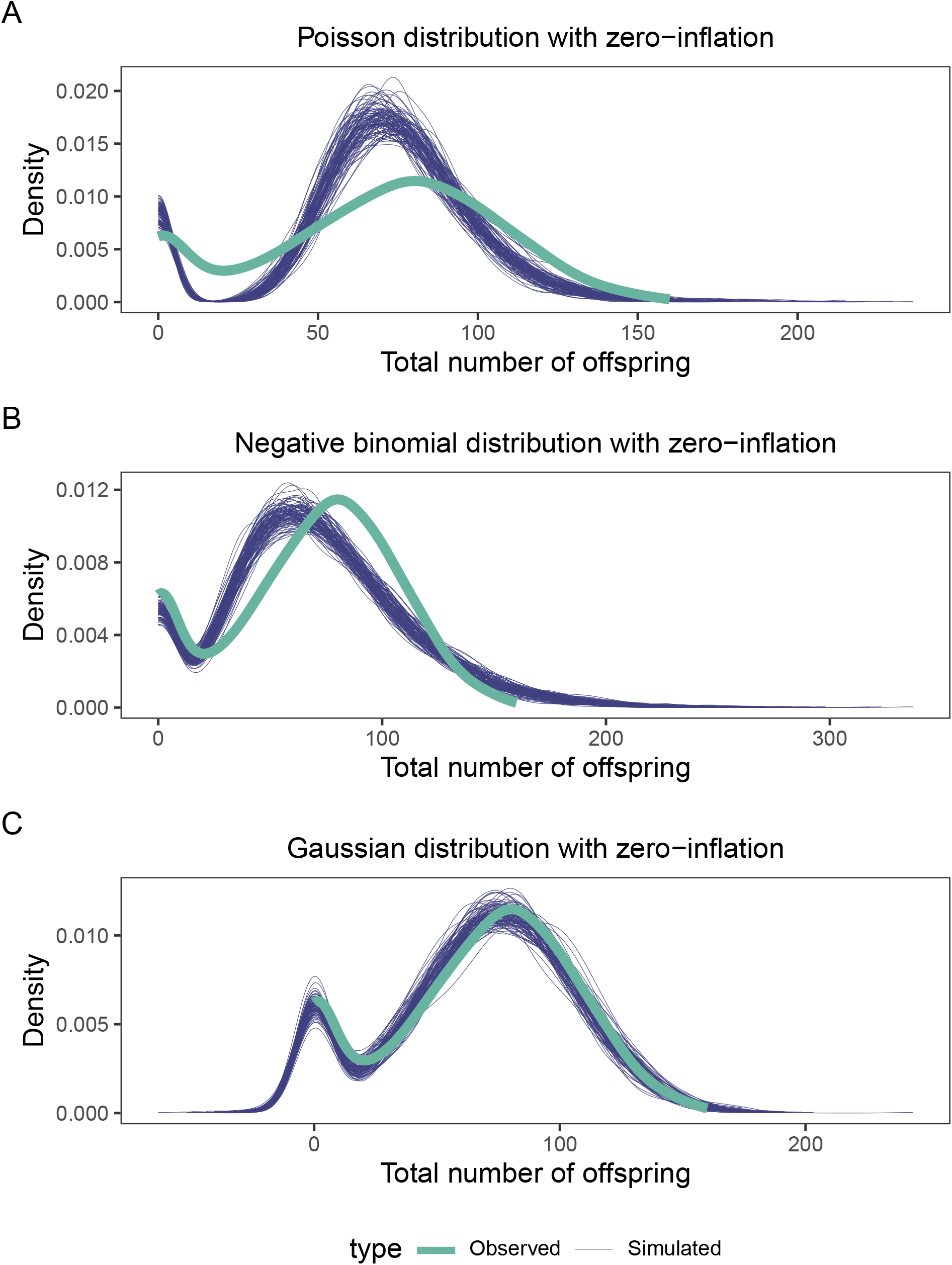
The distribution of the number of offspring in our data and those from 100 simulations of varying distributions. Simulations were done from a GLMM with a correction for zero-inflated data, and contains the full hierarchical model with different assumed distributions: **A)** Poisson, **B)** negative-binomial, and **C)** Gaussian.

**Table S2.**
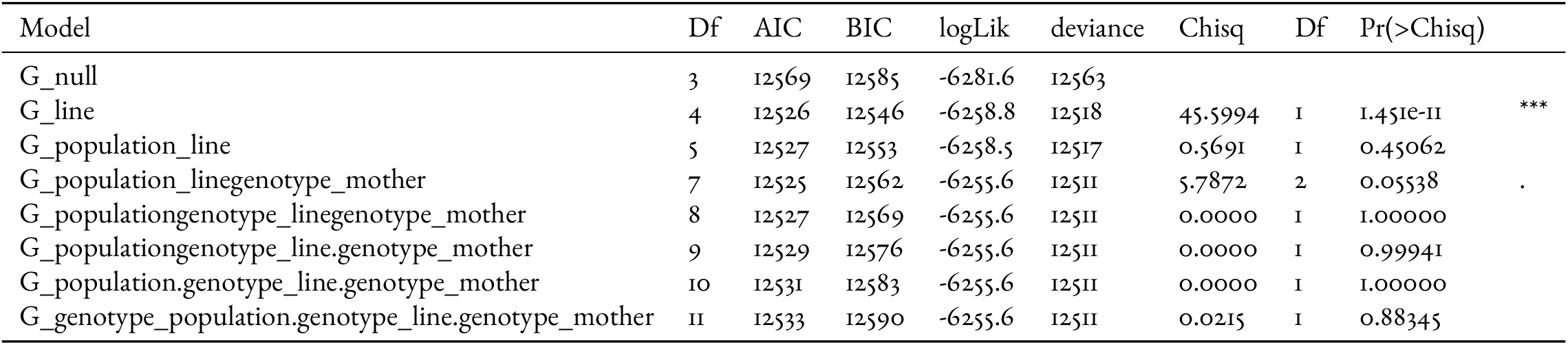
Model selection for total number of offspring. “G” stands for ‘Gaussian’. A dot between two terms means it includes that interaction in the random effect, whereas two concatenated terms means only an additive effect between the two terms in the random effect.

**Figure S12.**
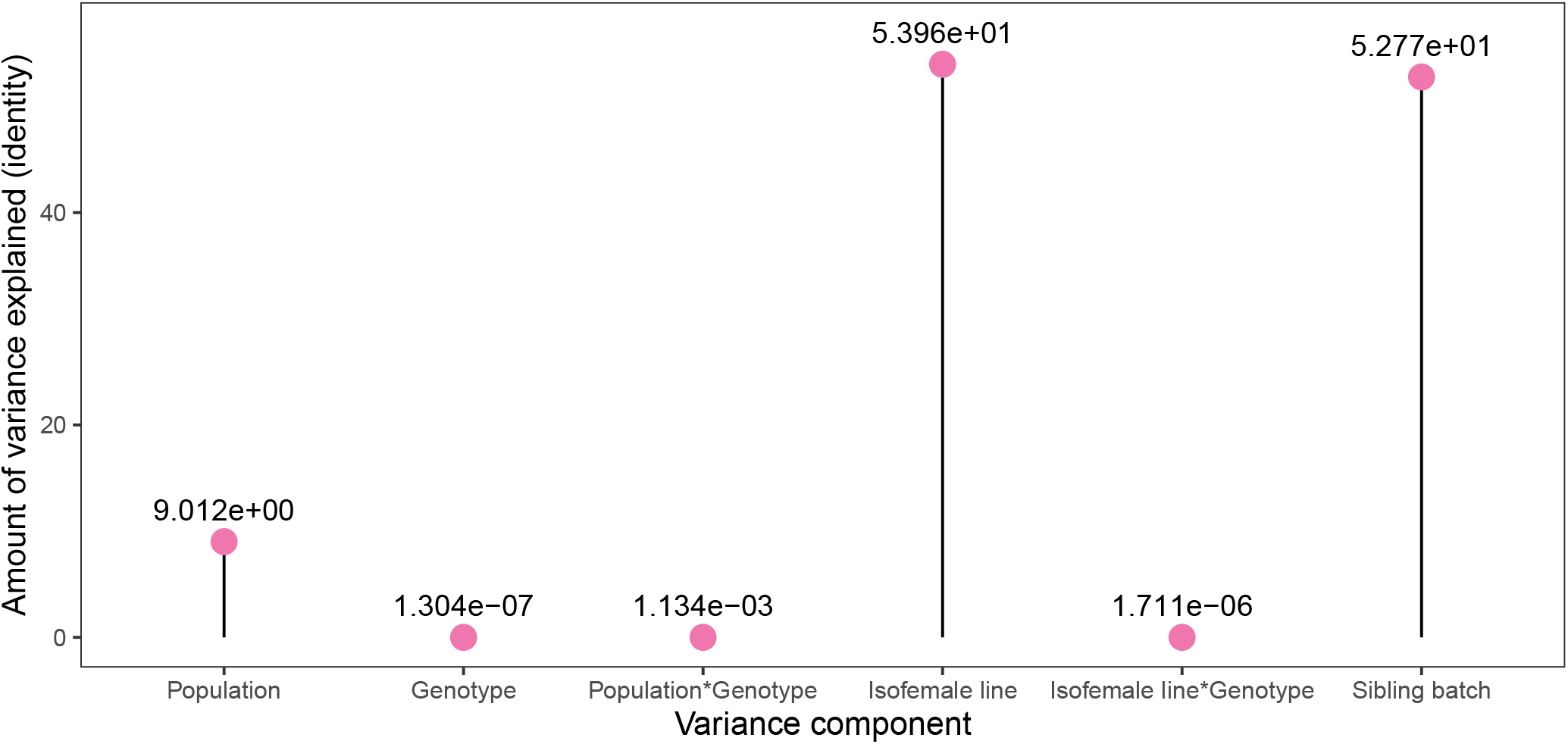
Variance in number of offspring partitioned to each hierarchical level of our experiment. These values are identity-linked.

**Figure S13.**
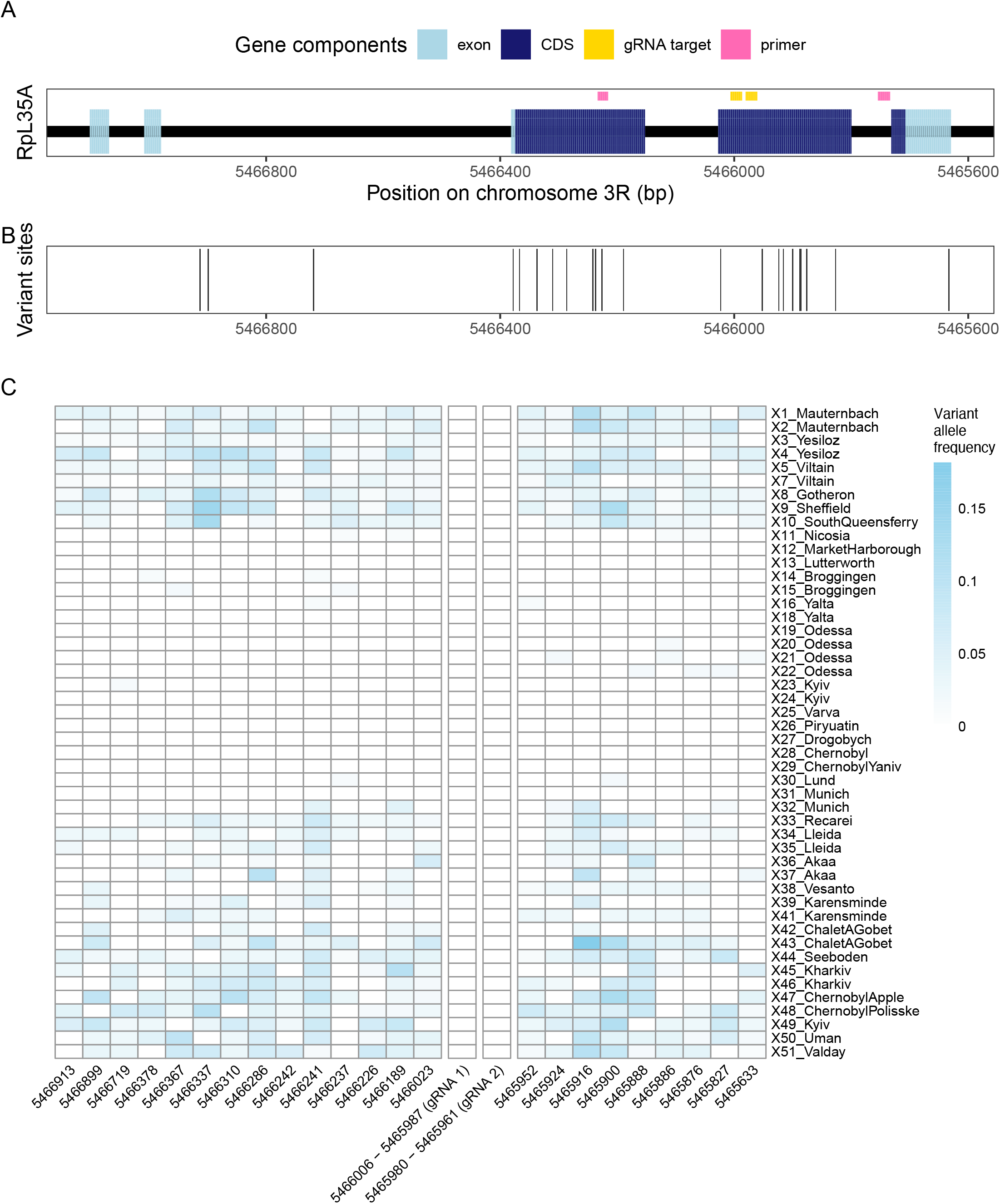
Variant sites at the *RpL35A* locus in the populations sampled by the DrosEU consortium. **A)** The *RpL35A* gene and its components, as well as the gene drive’s two gRNA target sequences and the two primers we used for amplification. **B)** Variant sites in the *RpL35A* gene. **C)** Allele frequencies of variant sites in the *RpL35A* gene in every population sampled by the DrosEU consortium (59), as well as the two gRNA target sequences (where there are no variants).

**Figure S14.**
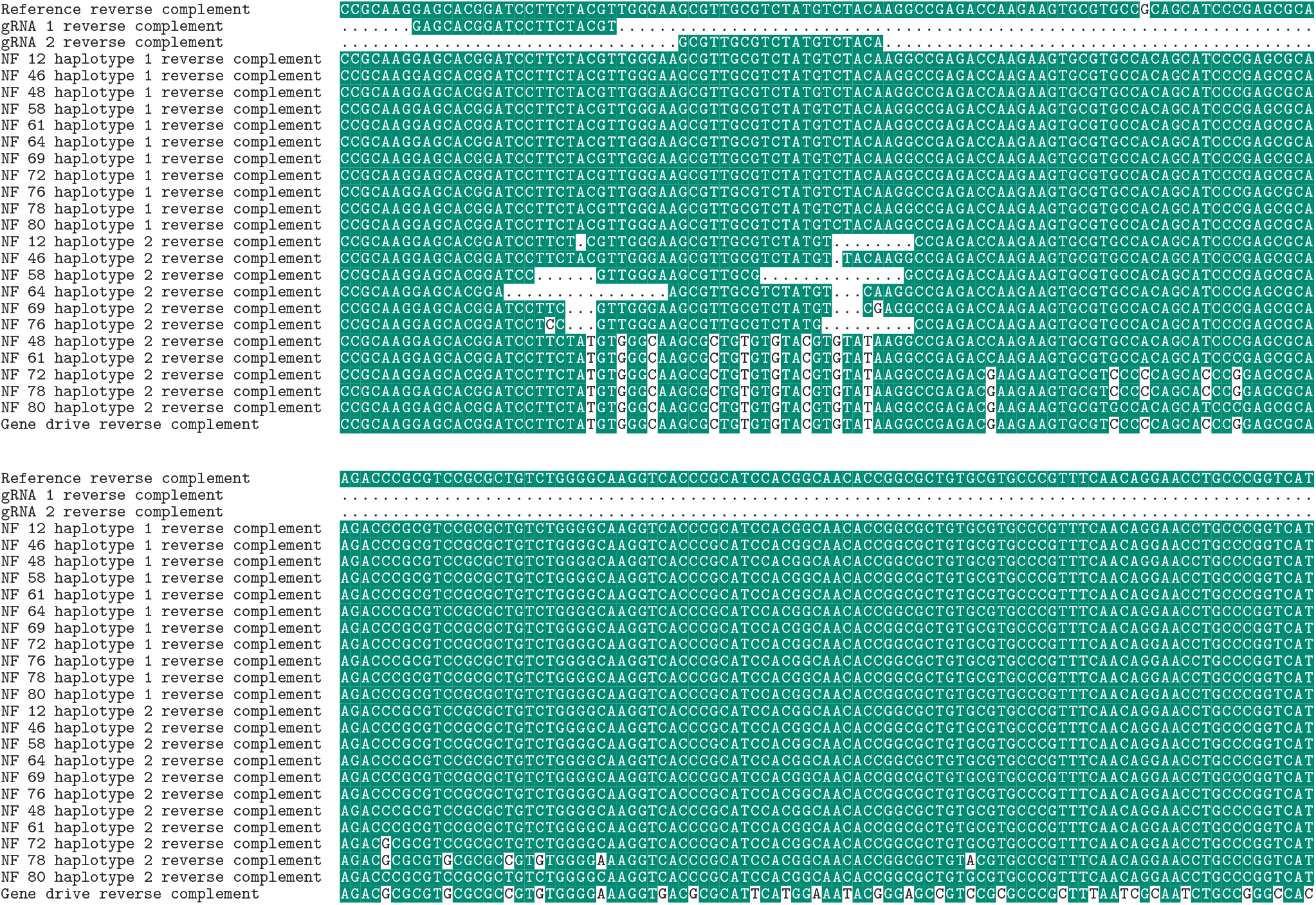
Multiple sequence alignment of the 11 observed RpL35A mutations. On top is the *D. melanogaster* reference genome, below which we show the gene drive’s two gRNA targets. Below the reference sequence, we show first the 11 wildtype haplotypes of each sample, followed by the 11 resistance haplotypes that were found, with white blocks indicating deletions and mismatches with the reference. At the bottom, we show the gene drive sequence. The order of the second haplotype sequences (the resistance alleles) is kept the same as in Figure 6. For the full alignment, see the GitHub repository (https://github.com/NickyFaber/HomingDrive_Heritability).

